# The Small GTPase Rab7 Regulates Release of Mitochondria in Extracellular Vesicles in Response to Lysosomal Dysfunction

**DOI:** 10.1101/2023.02.11.528148

**Authors:** Wenjing Liang, Rachel Y. Diao, Justin M. Quiles, Rita H. Najor, Liguo Chi, Benjamin P. Woodall, Leonardo J. Leon, Jason Duran, David M. Cauvi, Antonio De Maio, Eric D. Adler, Åsa B. Gustafsson

## Abstract

Mitochondrial quality control is critical for cardiac homeostasis as these organelles are responsible for generating most of the energy needed to sustain contraction. Dysfunctional mitochondria are normally degraded via intracellular degradation pathways that converge on the lysosome. Here, we identified an alternative mechanism to eliminate mitochondria when lysosomal function is compromised. We show that lysosomal inhibition leads to increased secretion of mitochondria in large extracellular vesicles (EVs). The EVs are produced in multivesicular bodies, and their release is independent of autophagy. Deletion of the small GTPase Rab7 in cells or adult mouse heart leads to increased secretion of EVs containing ubiquitinated cargos, including intact mitochondria. The secreted EVs are captured by macrophages without activating inflammation. Hearts from aged mice or Danon disease patients have increased levels of secreted EVs containing mitochondria indicating activation of vesicular release during cardiac pathophysiology. Overall, these findings establish that mitochondria are eliminated in large EVs through the endosomal pathway when lysosomal degradation is inhibited.

## Introduction

Cardiac myocyte contraction requires high levels of ATP which is supplied by mitochondrial oxidative phosphorylation. However, dysfunctional mitochondria generate excessive reactive oxygen species that can cause damage to cellular components, and these organelles can also directly activate cell death pathways^1^. Thus, efficient removal of aberrant mitochondria is crucial to maintain cellular homeostasis and cardiac function. To prevent unnecessary death, cells have evolved various mechanisms involved in repairing or removing dysfunctional mitochondria^2^. These cellular quality control pathways are particularly important in terminally differentiated cardiac myocytes, which are unable to dilute cellular damage through cell division and cannot be easily regenerated. Autophagy is the primary pathway involved in clearing dysfunctional mitochondria in the heart^3–5^ and involves the sequestration of a mitochondrion by an autophagosome which then delivers the cargo to a lysosome^2^. Mitochondria can also be engulfed by early endosomes^6, 7^, or directly taken up into lysosomes^8^. Although multiple mechanisms of mitochondrial degradation exist in cells, these pathways all converge at the lysosome which is ultimately responsible for the final breakdown of protein aggregates and organelles^9^. Whether alternative mitochondrial quality control pathways exist when lysosomal function is compromised or overwhelmed is currently unclear.

Cells are known to release vesicles from different origins that range from 0.05-1μm in diameter into the extracellular space^10^. Extracellular vesicles (EVs) are released after fusion of late endosomes (LE)/multivesicular bodies (MVB) with the plasma membrane or via budding of the plasma membrane. Studies have reported that these vesicles participate in cell communication by delivering nucleic acids, proteins, and lipids to recipient cells^10, 11^. The release of EVs is increased in various diseases or during stress. For instance, there is increased release of EVs from the heart during a myocardial infarction^12–14^. Secretion of EVs by skeletal muscle is also increased during exercise in both rodents and humans^15^.The EVs have been reported to promote regeneration of injured skeletal muscle tissue^16^. However, there is also emerging evidence that EVs might function as an alternative pathway of cellular quality control. EVs containing β-amyloid and α-synuclein have been detected in patients with Alzheimer’s disease and Parkinson’s disease, respectively^17, 18^. Cells have also been reported to release large (3.5-4μm) subcellular structures known as exophers which are much larger than traditional EVs^19, 20^. Intriguingly, several studies have identified mitochondrial proteins as EV cargo^20–23^, but their function and mechanism of release are still poorly understood. Formation and trafficking of endosomal vesicles containing EVs are regulated by Rab GTPases^10^. Rab7 is present on the LE/MVB and is required for the fusion between the LE/MVB and the lysosome^24^. It is also involved in regulating autophagosome-lysosome fusion^25, 26^, indicating crosstalk between the two internal degradation pathways. Whether *Rab7* plays a role in regulating fusion between the LE/MVB with the plasma membrane to release EVs into the extracellular space is still unknown.

Danon disease is caused by loss-of-function mutations in the lysosome-associated membrane protein 2 (*LAMP2*) gene, which leads to impaired autophagic degradation and development of cardiomyopathy^27, 28^. Interestingly, these patients are often asymptomatic in early childhood, suggesting that alternative mechanisms compensate for defective autophagic-lysosomal degradation. Here, we describe a new mechanism of mitochondrial clearance in cells that involves secretion of mitochondria in large EVs when lysosomal function is compromised. We report that the release of ubiquitinated proteins and mitochondria in large EVs from MVBs serves as an alternative cellular quality control pathway in the heart. *Rab7* functions as a switch in the trafficking of MVBs where its inactivation diverts trafficking of the LE/MVB to the plasma membrane for secretion of mitochondria-containing EVs. Overall, our findings suggest that this mechanism functions as an alternative cellular quality control pathway when lysosomal function is compromised to protect cells against accumulation of harmful cargo that is destined for degradation.

## Results

### Inhibiting lysosomal acidification leads to increased release of large and small extracellular vesicles

To investigate the relationship between lysosomal impairment and EV secretion, we treated mouse embryonic fibroblasts (MEFs) with the vacuolar H+ ATPase (V-ATPase) inhibitor Bafilomycin A1 (Baf A1) and examined EV release from these cells. Large and small EVs were collected from the conditioned medium using a differential centrifugation protocol^29^ (Supplementary Fig. 1a). Western blotting of EV fractions isolated from conditioned media showed that both large and small EV fractions were positive for Tsg101, Alix, and CD81, which are established markers of EVs derived from the MVBs^10^. These EV markers were significantly increased after Baf A1 treatment (Fig. 1a-b + Supplementary Fig. 1b), indicating increased secretion of EVs during inhibition of lysosomal acidification. Interestingly, the large EV fraction, but not small EVs, was also positive for the outer and inner mitochondrial membrane proteins Tom20 and Tim23 and treatment with Baf A1 significantly increased mitochondrial protein levels in large EVs (Fig. 1a-b).

**Figure 1.**
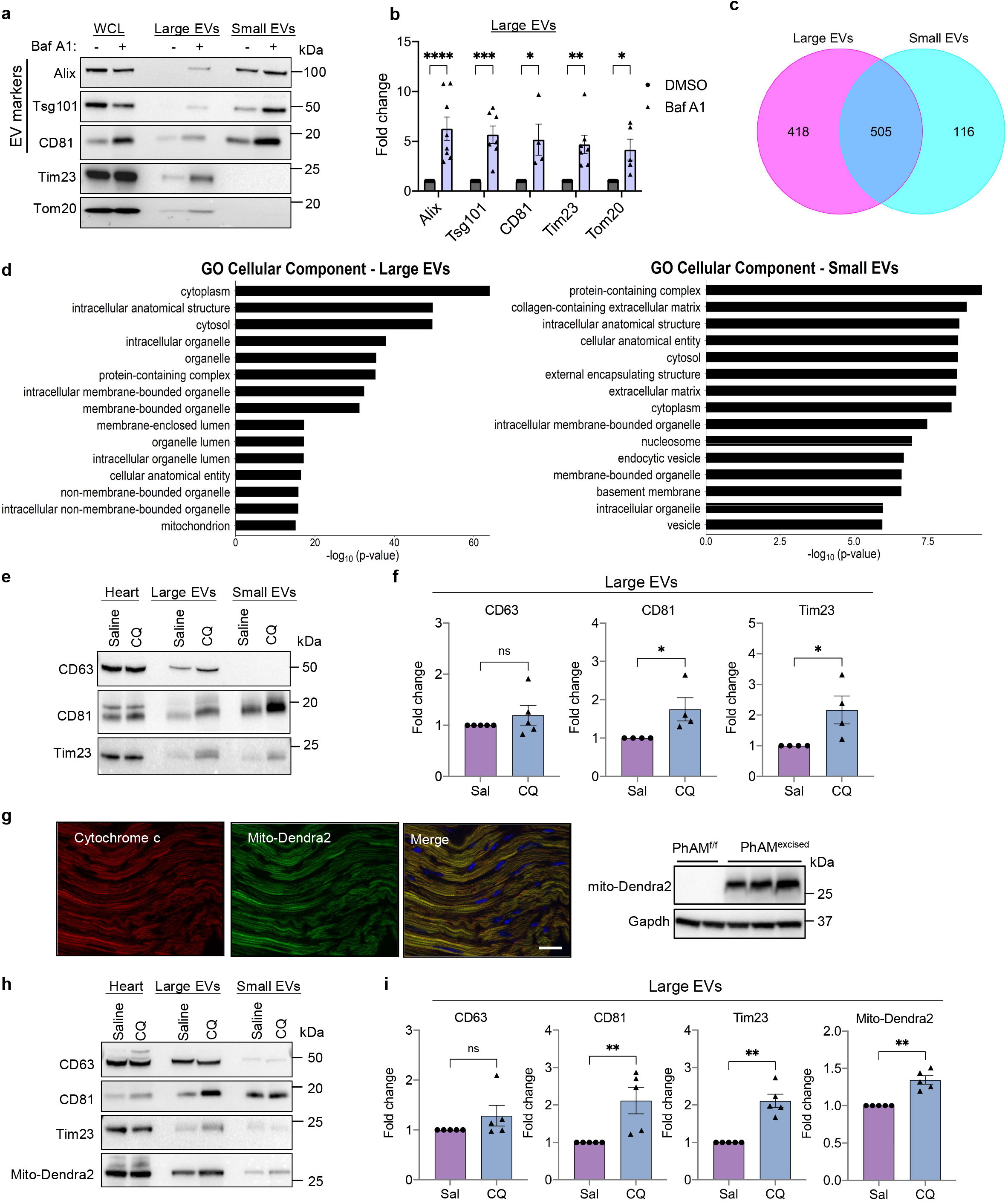
Inhibiting lysosomal acidification leads to increased release of large and small extracellular vesicles (EVs) from cells and heart. **a.** Representative Western blot analysis of whole cell lysate (WCL) and extracellular vesicle fractions for EV marker proteins Alix, Tsg101, CD81 and the mitochondrial proteins Tim23 and Tom20. The EVs were obtained by differential centrifugation of conditioned media from mouse embryonic fibroblasts collected over 48 h after treatment with vehicle or Bafilomycin (Baf A1, 50 nM). EVs from 1×10^7^ cells were loaded. **b.** Quantification of proteins in large EVs (n=4-8 biologically independent experiments). **c.** Venn diagram showing overlap of proteins detected in large and small EVs. **d.** Gene ontology (GO) analysis of the unique 418 proteins identified in large EV fractions with the top 15 terms for cellular component and biological processes plotted according to −log10False Discovery Rate (FDR). **e.** Representative Western blot analysis of EV proteins CD63, CD81 and the mitochondrial protein Tim23 in heart lysates and EV fractions isolated from heart tissue. **f.** Quantification of proteins in large EVs (n=4-5). **g.** PhAM^floxed^ mice were crossed with *Myh6-Cre* transgenic mice to generate a mouse line with cardiac specific expression of Mito-Dendra2. Expression of Mito-Dendra2 in myocytes was confirmed by imaging of frozen heart sections and by Western blotting of heart lysates. Scale bar=20μm. **h.** Representative Western blot analysis of CD63, CD81, Tim23, and Mito-Dendra2 in heart lysates and EV fractions isolated from heart tissue. **i.** Quantification of proteins in large EVs (n=5). Data are mean±SEM. *p<0.05,**p<0.01,***p<0.001 ****p<0.0001, ns = not significant.

Next, we performed proteomics analysis of large and small EVs to compare their composition and cargo. Among the proteins identified in small EVs, 81% (505/621) of these proteins were also present in large EVs (Fig. 1c). We also confirmed that proteins identified in large EVs overlapped with proteins in the exosomal and EV molecular databases Vesiclepedia (http://microvesicles.org/index.html) and Exocarta (www.exocarta.org) (Supplementary Fig. 1c). Importantly, none of the proteins identified were key autophagy proteins, suggesting that these vesicles are not representing autophagosomes. Furthermore, Gene Ontology (GO) enrichment analysis of the 418 proteins exclusively found in the large EVs showed an enrichment of membrane bound organelles including mitochondria (Fig.1d). In contrast, the small EVs showed an enrichment in proteins, cytosol and cytoplasm (Fig. 1d). Overall, these findings suggest that disrupting lysosomal acidification leads to increased secretion of large and small EVs from cells, that both originate from the MVBs in the endosomal pathway, and that large EVs may contain intact mitochondria.

To investigate if disrupting lysosomal function enhances the release of EVs *in vivo,*we administered chloroquine (CQ), a lysosomotropic drug that increases lysosomal pH, to mice and investigated changes in EV content within the extracellular space of heart tissue and in plasma isolated 24 h after administration (Supplementary Fig. 1d). While we observed no increase in circulating EVs in plasma after CQ administration (Supplementary Fig. 1e), we found that CQ treatment led to increased levels of large and small EVs in cardiac tissue (Fig. 1e-f + Supplementary Fig. 1f). Similar to what observed in MEFs, we also found increased levels of the mitochondrial protein Tim23 in the large EV fraction. To investigate whether the mitochondria in large EVs originated from cardiac myocytes, we generated mice with cardiac-specific expression of the Dendra2 fluorescent protein in mitochondria (mito-Dendra2) by breeding PhAM^f/f^ and Myh6-Cre mice (Fig. 1g). Large EV fractions isolated from hearts of mito-Dendra2 mice using differential centrifugation were positive for both Tim23 and mito-Dendra2 (Fig. 1h-i) suggesting that cardiac myocytes are a primary source of EVs released into the extracellular space. To validate our differential centrifugation protocol, we isolated EVs from cardiac tissue using bead conjugated antibodies against EV cell surface markers and these fractions were also positive for Tim23 and mito-Dendra2, and were significantly increased in mice treated with CQ (Supplementary Fig.1g). Together, these results suggest that mitochondria can be released in large EVs originating from the endosomal system when lysosomal function is compromised in the mouse heart.

### Rab7-deficiency leads to increased secretion of large and small EVs

Next, we investigated the mechanism by which the MVBs are directed to the plasma membrane for secretion of EVs rather than to lysosomes for degradation. Because the small GTPase *Rab7* facilitates fusion of the MVB with the lysosome^25, 26^, we investigated the role of *Rab7* in regulating MVB trafficking in cells. First, we evaluated the effect of Rab7-deficiency on the release of large and small EVs into conditioned media. Western blot analysis of EV fractions revealed that *Rab7*^-/-^ MEFs had increased release of large and small EVs into conditioned media at baseline compared to WT MEFs (Fig. 2a-b + Supplementary Fig. 2a). The large EVs were also positive for the mitochondrial protein Tim23 which increased only in the EV fractions from *Rab7*^-/-^ MEFs. Treatment of *Rab7*^-/-^ MEFs with GW4869, a known inhibitor of EV/exosome release^30^, diminished the release of mitochondria-containing large EVs from *Rab7*^-/-^ MEFs (Supplementary Fig. 2b). We further confirmed increased levels of entire mitochondria in the large EV fraction by quantifying mitochondrial DNA (mtDNA) by qPCR (Fig. 2c). Moreover, negative staining electron microscopy (EM) analysis demonstrated the typical cup-shaped morphology of the isolated small and large EVs (Fig. 2d). Furthermore, transmission electron microscopy (TEM) confirmed the presence of mitochondria in large EVs isolated from conditioned media of *Rab7*^-/-^ MEFs (Fig. 2e). Nanoparticle tracking analysis (NTA) showed a distinct size distribution of large EVs released by *Rab7*^-/-^ MEFs, while the size distribution of small EVs released by *Rab7*^-/-^ MEFs was similar to WT (Fig. 2f). Additionally, cargo proteins associated with large EVs secreted by WT and *Rab7*^-/-^ cells were identified by LC-MS/MS proteomics analysis. We identified 103 proteins that were significantly enriched in the EV fraction from *Rab7*^-/-^ MEFs and identified a variety of mitochondrial proteins (Fig. 2g). Similarly, Gene Ontology (GO) analyses also confirmed an enrichment of proteins associated with organelles and various metabolic processes in the cellular component and biological process categories, respectively, in EVs isolated from *Rab7*^-/-^ cells (Fig.2h). Taken together, these results suggest that Rab7-deficiency leads to increased release of large and small EVs from cells via the endosomal pathway. Analysis of various proteins involved in regulating endosome function and trafficking showed that CD81 and Alix protein levels were significantly increased in *Rab7*^-/-^ MEFs (Supplementary Fig. 2c). *Rab7*^-/-^ cells also had significant increased levels of Rab5 (early endosomes), Rab4 (recycling endosomes), Rab11 (recycling endosomes), and Rab9 (Golgi-derived vesicles) (Supplementary Fig. 2d). Arl8b, another small GTPase that resides on MVB and directs vesicle transport to the cell periphery^31^, was also significantly increased in Rab7-deficient cells (Supplementary Fig. 2d). Moreover, immunofluorescence analysis of CD81-positive vesicles in WT and *Rab7*^-/-^ MEFs confirmed increased co-localization between CD81-positive vesicles and mitochondrial Cytochrome *c* in *Rab7*^-/-^ MEFs at baseline and in WT cells after treatment with Baf A1 (Fig. 2i). The presence of mitochondria in CD81-positive vesicles in *Rab7*^-/-^ MEFs was also confirmed by correlative light and electron microscopy (Supplementary Fig.2e). Overall, these findings suggest that endosomal activity is enhanced in Rab7-deficient cells and that the Rab7-deficiency leads to redirection of mitochondria into CD81-positive vesicles for secretion.

**Figure 2.**
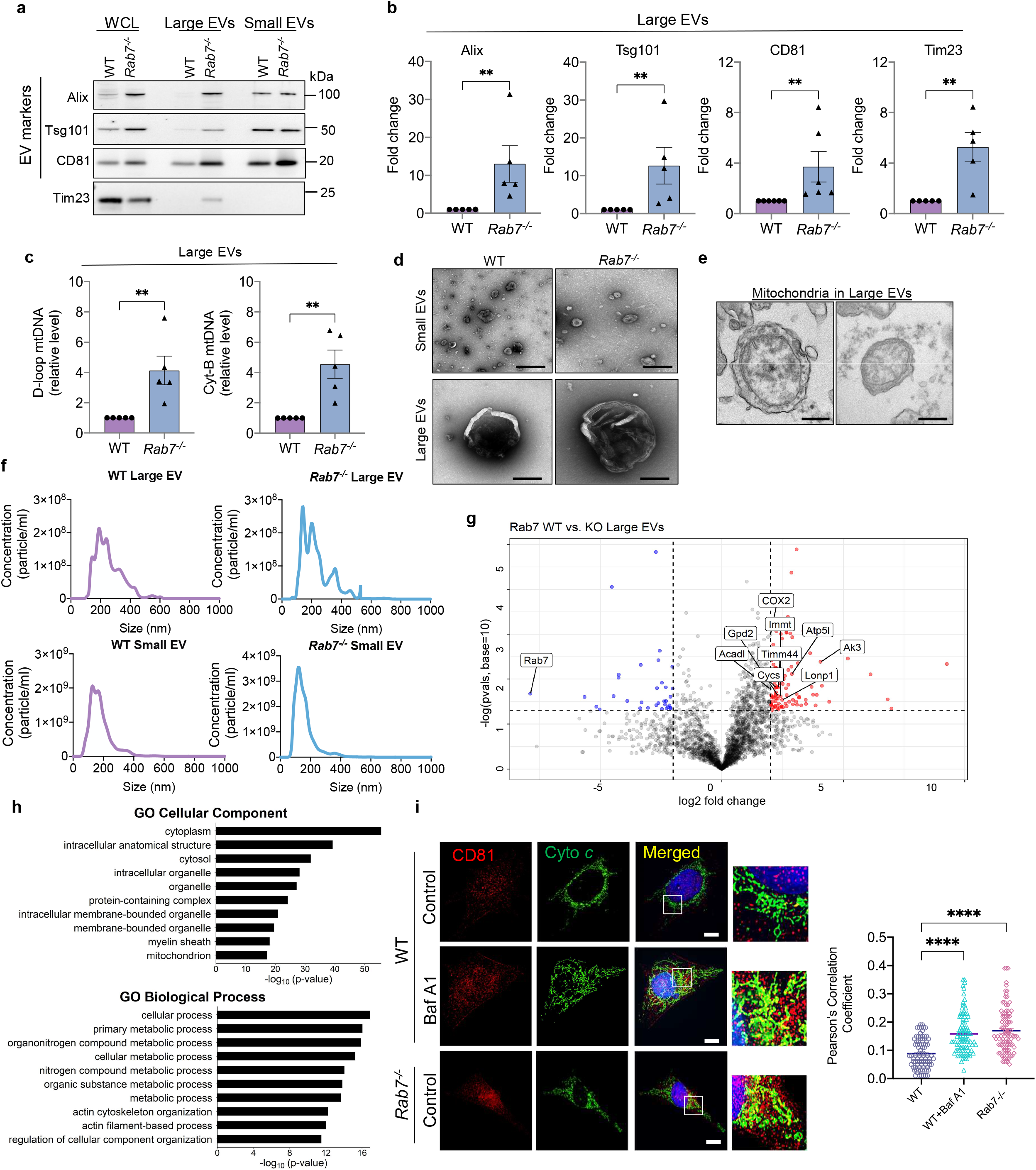
Rab7-deficient cells have increased EV secretion at baseline. **a.** Representative Western blot of Alix, Tsg101, CD81, and Tim23 in total cell lysates and EV preparations from WT and *Rab7*^-/-^ MEFs. **b.** Quantification of proteins in large EVs (n=5-6). **c.** Mitochondrial DNA (mtDNA) content in large EVs preparations (n=5). **d.** Visualization of small and large EVs isolated from conditional media were visualized by negative stain electron microscopy (scale bars=500 nm). **e.** Representative electron microscopy images of large EVs from *Rab7*^-/-^ MEFs containing mitochondria (scale bars=500 nm). **f.** Representative histograms of a Nanosight nanoparticle tracking analysis of large and small EVs secreted by wild type (WT) and *Rab7*^-/-^ MEFs (n = 4). **g.** Volcano plot of proteins identified proteins in large EV fractions from WT and *Rab7*^-/-^ large EVs. Proteins are plotted according to their −log10P values and log2 fold enrichment (WT/ *Rab7*^-/-^). Red dots represents proteins that are significantly enriched in *Rab7*^-/-^ EVs, while blue dots represents proteins that are decreased. **h.** Gene ontology (GO) enrichment analysis of EV proteins isolated from *Rab7*^-/-^ MEFs relative to wild type MEFs with top terms for cellular component and biological processes plotted according to −log10False Discovery Rate (FDR). **i.** Co-localization between CD81 and Cytochrome *c* in WT +/- Baf A1 (50nM for 24hr) and *Rab7*^-/-^ MEFs. Scale bar = 20μm. Pearson’s correlation coefficient (n=3 independent experiments with a total of 90 cells). Data are mean±SEM. **p<0.01, ****p<0.0001.

### Secretion of large EVs occurs independently of autophagy

Some studies have reported that secretion of certain vesicles from cells is dependent on autophagosome formation^20, 23^. Therefore, we examined whether secretion of mitochondria in large EVs required functional autophagy by evaluating EV release in WT and autophagy-deficient *Atg5*^-/-^ MEFs. Despite impaired autophagosome formation, the presence of Baf A1 caused a significant increase in the release of large EVs containing mitochondria *Atg5*^-/-^ MEFs (Fig. 3a-b). Small EV release was also increased by treatment with Baf A1 in *Atg5*^-/-^ MEFs (Supplementary Fig. 3a). In addition, mice with cardiac-specific impairment in autophagy due to deletion of *Atg7* had similar levels of myocardial EVs as WT mice (Fig. 3c-d + Supplementary Fig.3b). Interestingly, the EV fractions were also positive for LC3II, a protein presents on the autophagosome membrane^32^.

**Figure 3.**
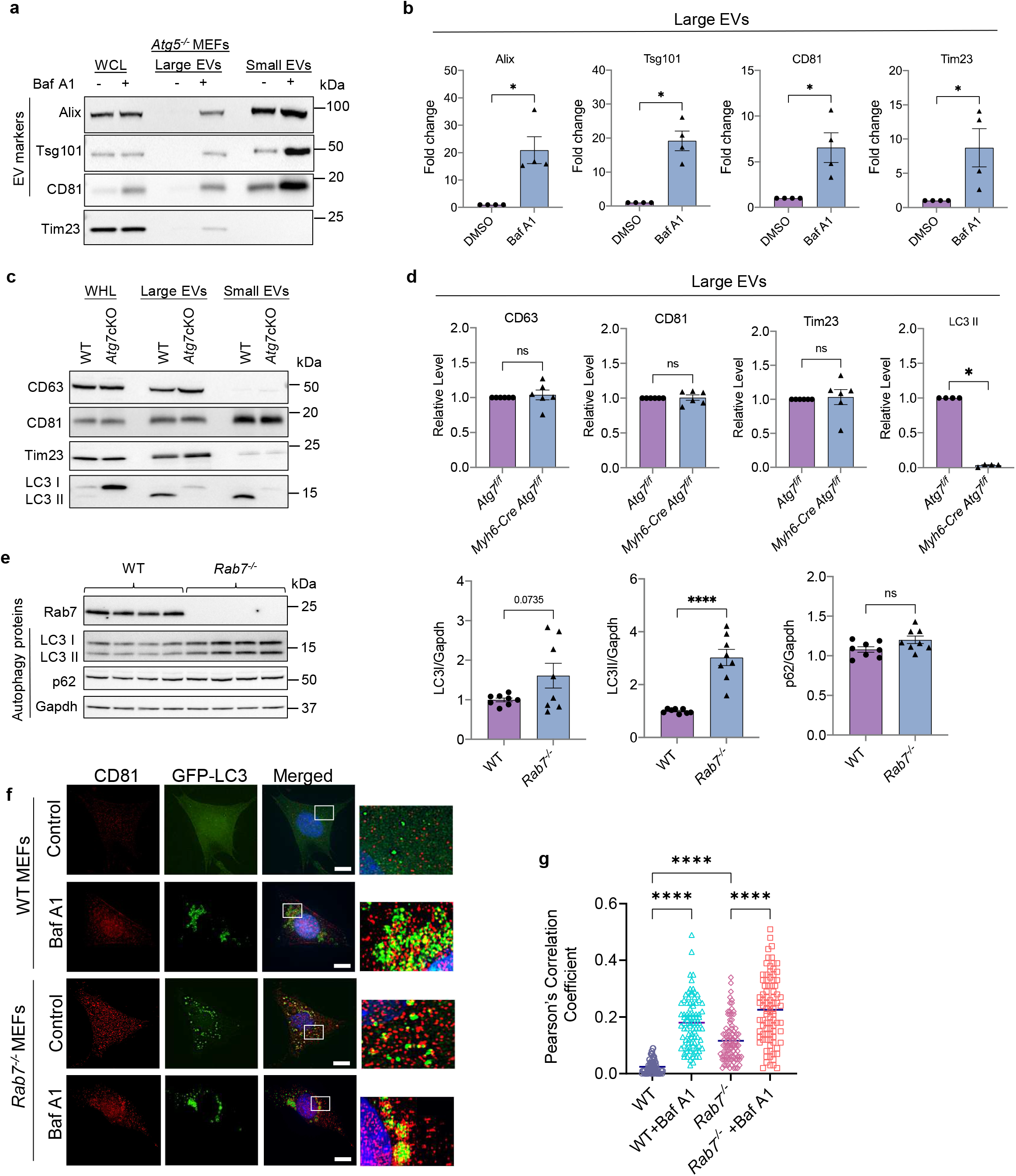
Secretion of large EVs is independent of the autophagy pathway. **a.** Representative Western blot of Alix, Tsg101, CD81, and Tim23 protein levels in total cell lysates and EV preparations from Atg5^-/-^ MEFs. The EVs were obtained by differential centrifugation of conditioned media from WT and *Atg5^-/-^* MEFs collected over 48 h after treatment with vehicle or Bafilomycin A1 (Baf A1, 50 nM). EVs from 1×10^7^ cells were loaded. **b.** Quantification of proteins in large EVs (n=4). **c.** Western blot analysis of CD63, CD81, Tim23 and LC3 protein levels in heart lysates and extracellular vesicle fractions isolated from heart tissue of *Atg7^f/f^* and *Myh6-Cre Atg7^f/f^* mice. **d**. Quantification of proteins in large EVs (n=4-6). **e.** Representative Western blot and quantification of LC3I, LC3II and p62 protein levels in WT and *Rab7*^-/-^ MEFs (n=8). **f.** Representative images of co-localization between CD81 and GFP-LC3 in WT and *Rab7*^-/-^ MEFs +/- Baf A1 (50nM for 24hr). **g.**Pearson’s correlation coefficient (n=3 independent experiments with a total of 90 cells scored). Scale bar = 20 μm. Data are mean±SEM. *p<0.05p<0.001 ****p<0.0001, ns = not significant.

Rab7 facilitates fusion between autophagosomes and lysosomes, and Western blot analysis confirmed that *Rab7*^-/-^ MEFs have significantly elevated levels of LC3II compared to WT MEFs (Fig. 3e), an indication of autophagosome accumulation. Unexpectedly, levels of the autophagy adaptor protein p62/SQSTM1 were unchanged in *Rab7*^-/-^ MEFs (Fig. 3e). Since p62 accumulates in cells when autophagy is impaired, unchanged p62 protein levels in *Rab7*^-/-^ MEFs suggests that an alternative mechanism of disposal might exist for p62-bound cargo. Autophagosomes have also been reported to fuse with endosomes to form hybrid structures known as amphisomes^33^. Therefore, we investigated whether disrupting lysosomal acidification or Rab7-deficiency induced fusion between autophagosomes and MVB. Immunofluorescence analysis showed increased co-localization between GFP-LC3 and CD81 in *Rab7*^-/-^ cells at baseline (Fig. 3f-g), indicative of amphisome formation^33^. The co-localization was also increased by Baf A1 treatment in WT and *Rab7*^-/-^ MEFs, suggesting a link between lysosomal dysfunction and amphisome formation. Taken together, these results demonstrate that secretion of large EVs is independent of autophagy. However, autophagosomes can fuse with MVBs to form amphisomes which might function to limit accumulation of autophagosomes containing cytotoxic cargo and allow for elimination of their cargo via secretion in EVs.

### Abrogation of EV secretion in *Rab7*^-/-^ MEFs enhances susceptibility to stress

Autophagy is important in clearing ubiquitinated cargo that has been labeled for degradation^2^. Despite the impairments in autophagy, we observed a small but significant decrease in ubiquitinated proteins in *Rab7*^-/-^ MEFs (Fig. 4a). However, we detected increased levels of ubiquitinated proteins in both large and small EVs secreted by *Rab7*^-/-^ MEFs (Fig. 4b), suggesting that the alternative vesicular pathway functions to dispose ubiquitinated cargo that would otherwise be degraded.

**Figure 4.**
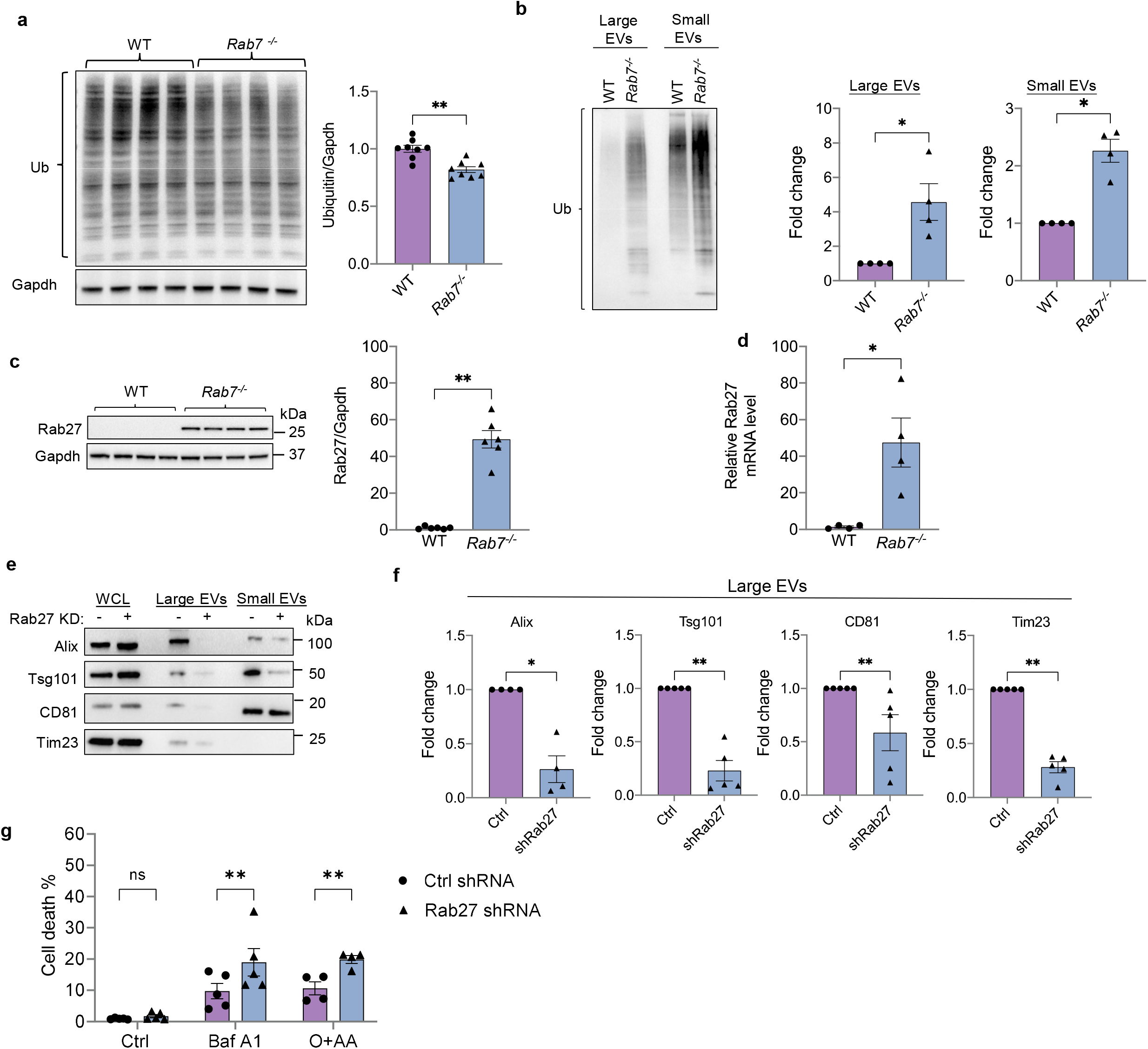
Rab27 knockdown in Rab7-deficient cells abrogates secretion of large EVs containing mitochondria and increases susceptibility to stress. **a.** Representative Western blot and quantification of ubiquitinated proteins in WT and *Rab7*^-/-^ MEFs at baseline (n=8). **b.** Representative Western blot and quantification of total ubiquitin levels in small and large EVs secreted by WT and *Rab7*^-/-^ MEFs at baseline (n=4). **c.** Representative Western blot and quantification of Rab27 protein levels in WT and *Rab7*^-/-^ MEFs (n=6). **d.** Rab27 transcript levels in WT and *Rab7*^-/-^ MEFs (n=4). **e.** Western blot of Alix, Tsg101, CD81, and Tim23 protein levels in cell lysates and EV preparations from *Rab7*^-/-^ MEFs infected with control or Rab27a shRNA. **f.** Quantification of proteins in large EVs (n=5-6 independent experiments). **g.** Analysis of cell death using Yo-Pro1 in *Rab7*^-/-^ MEFs infected with control or Rab27a shRNA. Cells were treated with Baf A1 (50nM) for 24h or O+AA (10μM+10μM) for 24h (n=4-5). Data are mean±SEM. *p<0.05, **p<0.01, n.s. not significant.

Rab27 is an important regulator of EV secretion and facilitates docking of MVBs to the plasma membrane^34^. Strikingly, we discovered that Rab27 protein and mRNA levels were significantly increased ~45-fold in *Rab7*^-/-^ MEFs compared to WT cells (Fig. 4c-d). Knockdown of Rab27 in *Rab7*^-/-^ MEFs abrogated the release of both large and small EVs (Fig. 4e-f + Supplementary Fig. 4a-b), and was associated with a small but significant increase in levels of ubiquitinated proteins (Supplementary Fig. 4c), supporting the notion that EVs release facilitates elimination of ubiquitinated cargo from cells. Finally, to examine if inhibiting EV release in *Rab7*^-/-^ MEFs alters susceptibility to stress, we treated cells with Baf A1 to disrupt lysosomal acidification or with Oligomycin+Antimycin A (O+AA) to inhibit mitochondrial respiration. Loss of Rab27-mediated EV release exacerbated cell death in *Rab7*^-/-^ MEFs following either treatment (Fig. 4g), suggesting that this is an important mechanism in limiting accumulation of cytotoxic material in cells.

### Genetic ablation of *Rab7* in the adult heart leads to increased secretion of EVs into cardiac tissue

To investigate the role of Rab7 in regulating EV release *in vivo*, we crossed *Rab7^f/f^* mice with the tamoxifen-inducible Myh6-MerCreMer (*MCM*) transgenic mice to allow for myocyte-specific deletion of *Rab7* in the adult heart^35^. *MCM*, *Rab7^f/f^* and *Rab7^f/f^ MCM* mice were injected with 5 doses of tamoxifen and cardiac function was evaluated 28 days later (Supplementary Fig. 5a). Echocardiographic assessment revealed no significant changes in left ventricular contractile function or structure (Fig. 5a + Supplementary Fig. 5b). Gross examination of hearts revealed a small but significant increase in heart weight to body weight (HW/BW) ratio in *Rab7^f/f^ MCM* mice compared to *MCM and Rab7^f/f^* mice (Fig. 5b), suggesting development of mild cardiac hypertrophy. However, histological analysis of hearts demonstrated normal cardiac structure with no fibrosis in *Rab7^f/f^* MCM mice (Fig. 5c). Mitochondrial respiration was also normal in *Rab7^f/f^ MCM* hearts (Supplementary Fig. 5c). Evaluation of changes at the ultrastructural level by TEM confirmed normal myofibrillar structure and mitochondria but revealed the presence of many small vacuoles in Rab7-deficient myocytes (Fig. 5d).

**Figure 5.**
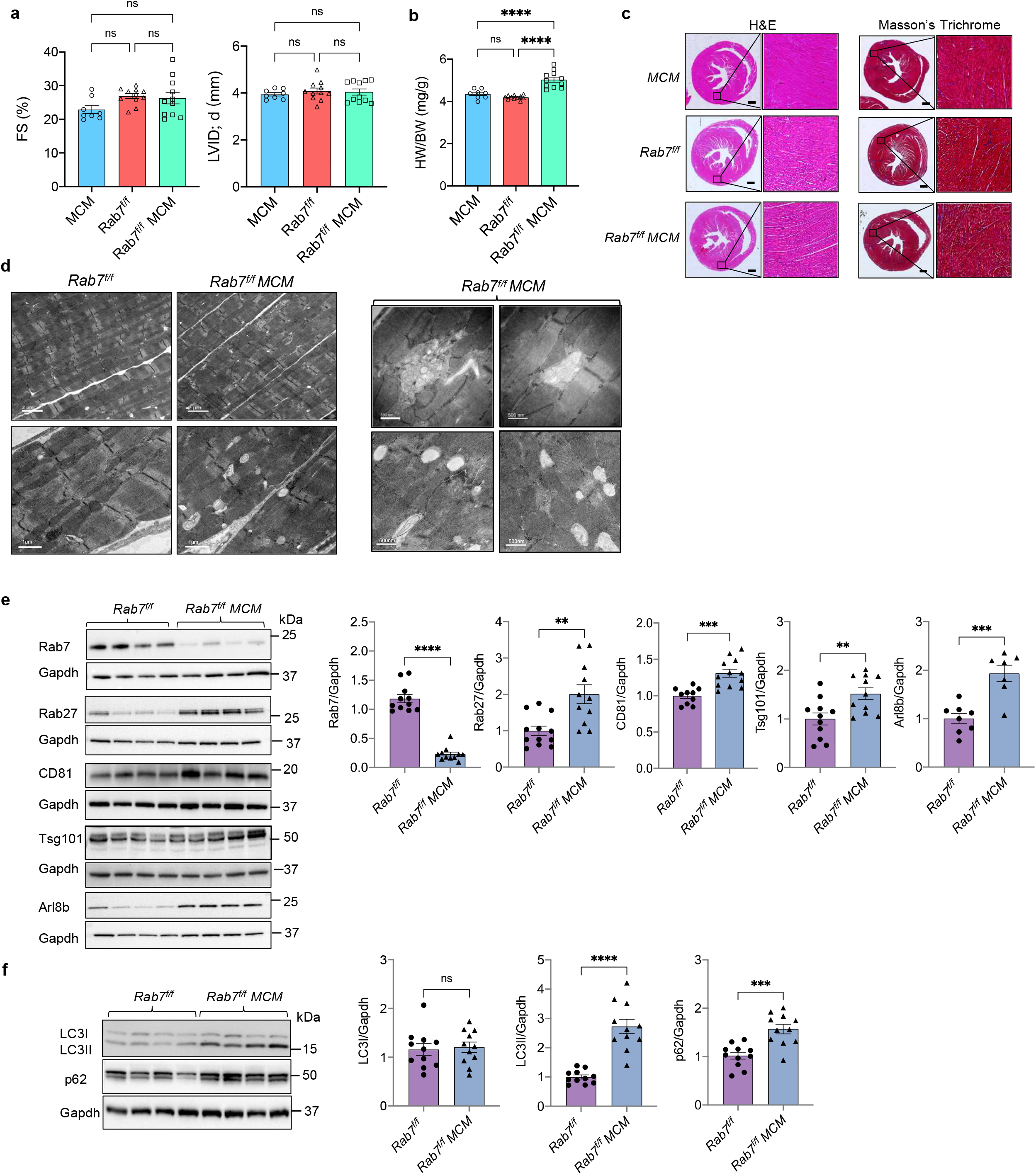
Characterization of mice with cardiac specific deletion of *Rab7*. **a.** Echocardiographic analysis of ventricular function and structure at 28 days post-tamoxifen treatment. Percent (%) fractional shortening (FS) and left ventricular end-diastolic dimension in diastole (LVID; d). *MCM* (n=8), *Rab7^f/f^* (n=11), *Rab7^f/f^ MCM* (n=11). **b**. Heart weight to body weight ratio (HW/BW), n=8-11 per group). **c.** H&E and Masson’s Trichrome staining of heart sections. Scale bar=0.5mm. **d.** Ultrastructural analysis by transmission electron microscopy at D28. **e.** Representative Western blots and quantification of endosomal proteins (n=7-12). **f.** Representative Western blots and quantification of autophagy proteins (n=11). Data are mean±SEM. **p<0.01, ***p<0.001, ****p<0.0001, ns = not significant.

Next, we examined whether loss of Rab7 in the heart was associated with changes in proteins involved in regulating endosomal trafficking and EV secretion. We found that Rab27, CD81, Tsg101 and Arl8b were significantly increased in Rab7-deficient hearts (Fig. 5e). Because Rab7 functions in the autophagy pathway, we also examined the effect of *Rab7* deletion on autophagic activity in hearts. Consistent with our *in vitro* findings, autophagic flux was impaired in Rab7-deficient hearts as assessed by significant accumulation of LC3II and the autophagy adaptor protein p62 (Fig. 5f).

Because mitochondrial respiration and cardiac function were normal in Rab7-deficient hearts despite impaired autophagy, we investigated whether loss of Rab7 in the heart led to enhanced secretion of EVs. Indeed, Rab7-deficienct hearts had increased levels of large EVs in the cardiac tissue (Fig. 6a-b+ Supplementary Fig. 6a), whereas *MCM* mice had similar EV release as WT (Supplementary Fig. 6b-c). Negative staining EM confirmed the cup-shape morphology of large EVs isolated from the *Rab7^f/f^* and *Rab7^f/f^ MCM* heart tissue (Fig. 6c) while NTA showed that vesicle sizes ranged from ~120nm to 780nm (Supplementary Fig. 6d). We also found increased levels of the mitochondrial protein Tim23 as well as elevated mtDNA content in large EVs isolated from *Rab7^f/f^ MCM* hearts (Fig. 6a, b, and d). To further confirm that the mitochondria detected in EVs originated from cardiac myocytes, we isolated total EVs (large and small) from cardiac tissue of *Rab7^f/f^*; *Mito-Dendra2* and *Rab7^f/f^ MCM*; *Mito-Dendra2* mice using bead conjugated antibodies against EV cell surface markers. Western blot analysis of EVs isolated from hearts of *Rab7^f/f^ MCM*; *Mito-Dendra2* mice showed that they were positive for both Tim23 and mito-Dendra2 (Fig. 6e-f), confirming that the mitochondria originated from cardiac myocytes. Altogether, these results demonstrate that cardiac specific Rab7-deficiency leads to impaired autophagic degradation concurrent with increased release of large EVs containing mitochondria in the heart.

**Figure 6.**
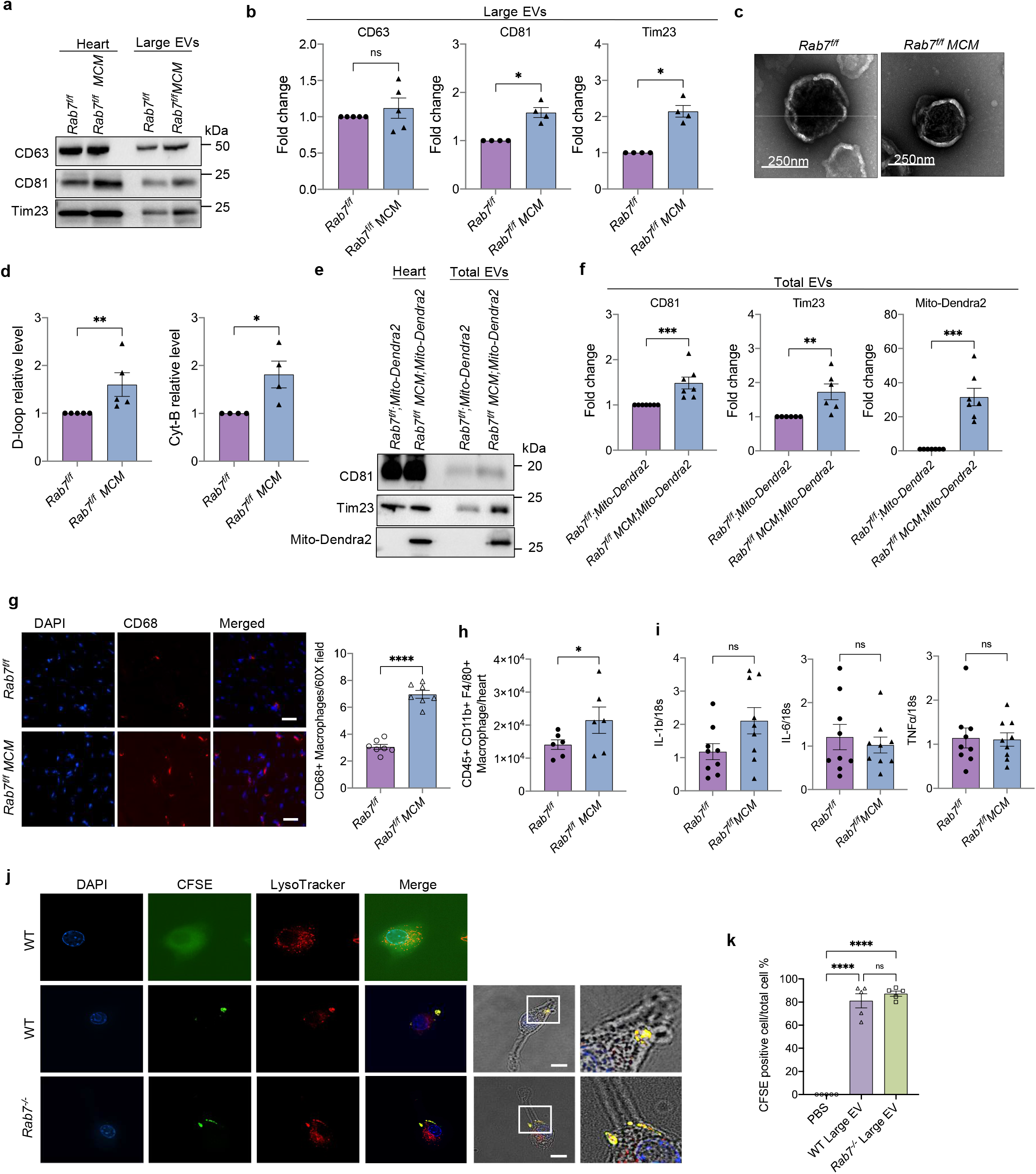
Enhanced secretion of large EVs containing mitochondria in Rab7-deficient hearts. **a.** Representative Western blot of CD63, CD81 and Tim23 in whole heart lysates and extracellular vesicle fractions from *Rab7^f/f^* and *Rab7^f/f^ MCM* heart tissue. **b.** Quantification of protein levels in large EV fractions (n=4-5) **c.** Negative stain electron microscopy of large EVs isolated from *Rab7^f/f^* and *Rab7^f/f^* MCM heart tissue. **d.** Mitochondrial DNA (mtDNA) content in large EVs preparations (n=4-5). **e.** Representative Western blot of CD81, Tim23, and Mito-Dendra 2 protein levels in whole heart lysates and extracellular vesicle fractions from *Rab7^f/f^*;*Mito-Dendra2* and *Rab7^f/f^MCM*;*Mito-Dendra2* heart tissue. Large and small (total) EVs were isolated from hearts using immunoaffinity capture method. **f.** Quantification of protein levels in the EV fractions (n=6-7). **g.** Immunostaining and quantification of CD68-positive cells in *Rab7^f/f^* and *Rab7^f/f^ MCM* hearts (n=7), Scale bar=20μm. **h.** Cardiac macrophage content in *Rab7^f/f^* and *Rab7^f/f^ MCM* hearts analyzed by flow cytometry (n=6). **i.** qPCR analysis for inflammatory markers in *Rab7^f/f^* and *Rab7^f/f^ MCM* hearts (n=9). **j.** Assessment of large EV uptake by RAW 264.7 macrophages. Representative images of cells after incubation (24h) with CFSE-labelled EVs (green). Blue (dapi) and red (Lysotracker Red). Scale bar=10μm. **k.** Quantification of CFSE-labeled EV uptake by cells (n=5, a minimum of 100 cells were scored in each independent experiment). Data are mean±SEM. *p<0.05, **p<0.01, ***p<0.001, ****p<0.0001, ns = not significant.

Similar to CQ treatment of WT mice, we observed no increase in circulating EVs in plasma after deletion of *Rab7* in myocytes (Supplementary Fig. 6e), suggesting that the secreted EVs remain in the cardiac tissue. Therefore, we investigated the fate of released EVs in *Rab7^f/f^ MCM* mice. The primary function of macrophages is to dispose unwanted material through phagocytosis^36^. Immunofluorescent staining of heart sections and flow cytometry showed that Rab7-deficient hearts displayed a significant increase in cardiac macrophages (Fig. 6g-h + Supplementary Fig. 6f), however, no transcriptional changes were observed in inflammatory markers (Fig. 6i). To assess if macrophages take up large EVs, carboxyfluorescein succinimidyl ester (CFSE)-labelled EVs isolated from conditioned media of WT or *Rab7*^-/-^ MEFs were added to cultured RAW 264.7 macrophages. After 24 h of incubation, a majority of cells were positive for CFSE (Fig. 6j-k), and the CFSE fluorophore co-localized with LysoTracker Red, indicating delivery of EVs to lysosomes after uptake. Therefore, loss of Rab7 in adult myocytes amplifies EV release during autophagy dysfunction and this response is sufficient to maintain cardiac homeostasis in vivo as cardiac macrophages ensure phagocytosis of released cargo.

### Increased secretion of EVs in hearts with age and in Danon Disease

The burden on lysosomes to eliminate damaged proteins and organelles increases with age in long-lived cells such as cardiac myocytes. Studies have reported various age-related changes in lysosomes ranging from alterations in size and number to impaired lysosomal hydrolase activity^37^. To investigate if aging affects the release of EVs *in vivo*, we evaluated the level of EV secretion in hearts of young (4-month-old) and aged (24-month-old) WT mice. We observed a significant increase in levels of large EVs containing mitochondria in cardiac tissue from aged mice using differential centrifugation (Fig. 7a-b) or bead conjugated antibodies against EV cell surface markers (Fig. 7c). A loss-of-function mutation in *LAMP2* is known to underlie Danon disease which is associated with a defect in autophagic-lysosomal-mediated degradation. To investigate if Lamp2-deficiency in mice correlated with increased EV secretion, we isolated large EVs from hearts of WT and *Lamp2*^-/-^ mice at 4 months of age, prior to any cardiac dysfunction^27, 38^. Western blot analysis showed increased levels of large EVs containing mitochondria in *Lamp2*^-/-^ hearts (fig.7d-e). Finally, we analyzed EV content in cardiac biopsies obtained at the time of explant from two Danon Disease patients, a young male (17 years) who underwent total artificial heart implantation and a female (39 years) who underwent heart transplantation. Immunoaffinity capture of EVs (large and small) using bead conjugated antibodies against EV cell surface markers showed that the Danon Disease patients had higher levels of EVs positive for mitochondria in cardiac tissue compared to normal control hearts (Fig. 7f). Collectively, these results confirm that a compromise in internal degradation pathways due to age or genetic mutations leads to increased secretion of mitochondria-containing EVs from cells.

**Figure 7.**
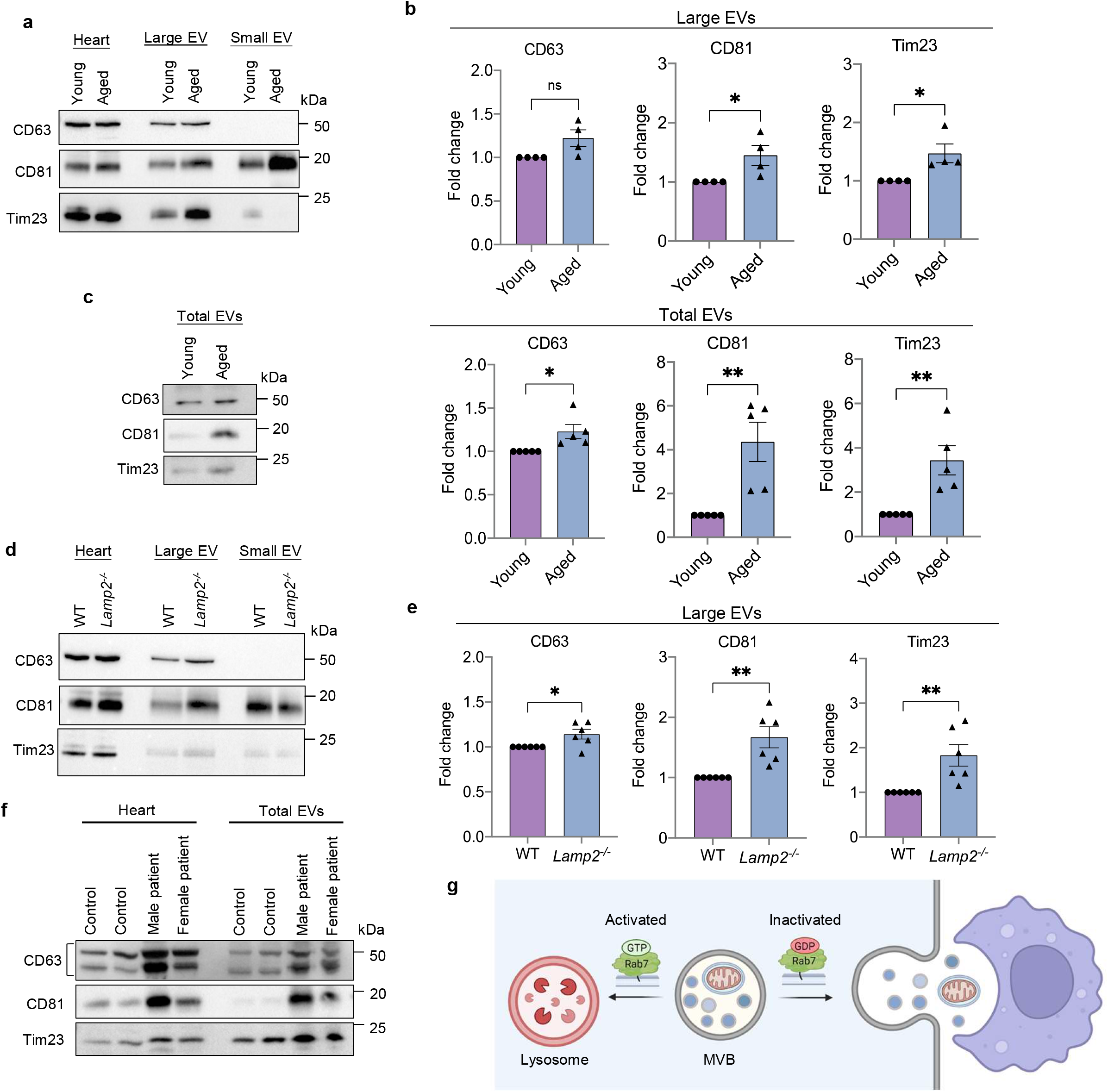
Enhanced secretion of large EVs containing mitochondria in aged and Lamp2-deficient hearts. **a.** Representative Western blot of CD63, CD81, and Tim23 in whole heart lysates and cardiac tissue EVs prepared from young (4 month) and aged (24 month) mice using differential centrifugation. **b.** Quantification of proteins in large EV fractions (n=4). **c.** Representative Western blot and protein quantification of total (large and small) EVs in hearts using immunoaffinity capture (n=5). **d.** Representative Western blot of CD63, CD81 and Tim23 in whole heart lysates and cardiac tissue EVs prepared from WT and *Lamp2*^-/-^ mice using differential centrifugation. **e.** Quantification of proteins in large EV fractions (n=6). **f.** Western blot of heart lysates and cardiac tissue extracellular vesicles (EVs) prepared from control and Danon Disease patients (male and female). **g.** Overview of Rab7 activation and EV secretion in cells. Data are mean±SEM. *p<0.05, **p<0.01, ns = not significant.

## Discussion

Mitochondria play a central role in cardiac energy metabolism and are responsible for generating most of the energy needed to sustain contraction. Because cardiac myocytes are post-mitotic, maintaining a healthy population of mitochondria is essential for their long-term survival. Intracellular degradation pathways that are responsible for degrading defective and potentially harmful mitochondria rely on lysosomes for the breakdown of cargo. In this study, we identified an alternative pathway for elimination of mitochondria and ubiquitinated proteins from cells when lysosomal function is compromised that involves secretion of mitochondria in large EVs. The EVs originate in late endosomal compartments that fuse with the plasma membrane to release the intralumenal vesicles into to the extracellular environment (Figure 7g). Although this pathway is independent of autophagy, autophagosomes can deliver cargo to MVB fusion to form an amphisome. Our findings also demonstrate that the small GTPase Rab7 functions as a switch in the trafficking pathway and that inactivation of Rab7 leads to transport of the MVB to the plasma membrane. The secreted large EVs do not enter the circulation to a significant extent but undergo phagocytosis by adjacent macrophages for delivery to lysosomes. Overall, these findings demonstrate that secretion of mitochondria in EVs can serve as an alternative pathway of eliminating cellular components when internal degradation pathways are compromised.

It is well established that the internal pathways responsible for eliminating plasma membrane receptors, protein aggregates or organelles rely on lysosomes for degradation and recycling of cargo^9^. However, lysosomal function declines with age^37^ and acidification can be impaired by the chemotherapeutic drug doxorubicin^39^. In addition, loss-of-function mutations in the lysosomal protein *LAMP2* causes Danon Disease, which leads to profound cardiac hypertrophy and heart failure^28^. Although the penetrance of disease is nearly 100% in males, affected individuals are often asymptomatic in early childhood. Similarly, *Lamp2*^-/-^ mice accumulate autophagosomes due to impaired flux, but do not display left ventricular dysfunction until a later age^38^. This suggests that an alternative mechanism exists in the heart and that other tisssues may temporarily compensate for the defect in lysosomal function. Accordingly, our findings demonstrate that disrupting lysosomal function pharmacologically or genetically in cells and mice led to increased secretion of both small and large EVs from cells. More importantly, conditional deletion of Rab7 in the heart had little effect on mitochondrial health and cardiac function despite compromised autophagy and accumulation of autophagosomes. Instead, there is an increase in the release of EVs into the extracellualar space of these hearts, suggesting that this functions as an alternative cellular quality control pathway that allows the cell to eliminate cytotoxic materials. Increased levels of secreted EVs were also identified in hearts of Lamp2-deficient mice and in pateints with Danon Disease.

Cells can secrete many types of vesicles of various sizes and origins. For instance, adult neurons and cardiac myoyctes can extrude protein aggregates and mitochondria in large (3.5-4μm) vesicular structures called exophers^19, 20^. Their formation is linked to autophagy where suppression of autophagy reduces exopher formation while activation of autophagy increases exopher generation in cardiac myocytes^20^. Similalry, Phinney and colleagues reported that mesenchymal stem cells can transfer mitochondria to macrophages by secretion in microvesicles^23^. The authors observed that mitochondria sequested in autophagosomes are transported to the plasma membrane, where they are released by outward budding of the plasma membrane. In contrast, we showed that the large EVs (~120-800nm) secreted from cells were similar to small EVs (i.e. exosomes) and originate from the same endosomal pathway. Their formation was independent of the autophagy process. However, autophagosomes can clearly utilize this secretion pathway to eliminate cargo from cells by fusing with endosomes to form hybrid organelle structures known as amphisomes^33^. Although the amphisome is thought to represent a transitional state prior to fusion with a lysosome, there is also evidence that this hybrid vesicle can fuse with the plasma membrane to release cargo^33^. Thus, it is possible that the fusion with MVB leads to gain of proteins involved in facilitating transport of the hybrid vesicle to the plasma membrane for fusion. Excessive accumulation of autophagosomes due to compromised fusion between autophagosomes and lysosomes can be cytotoxic and lead to cardiomyopathy^40^. Thus, formation of the amphisome could represent a mechansism to facilitate secretion of cargo to limit autophagosome accumulation and redcue cardiotoxicity.

It is well established that EVs can function as signaling vesicles by delivering nucleotides and proteins to other cells^10^. Uptake of encapsulated mitochondria has also been reported to influence cellular function after uptake. For instance, microvesicles enriched in mitochondria released by monocytic cells are pro-inflammatory and induce type I Interferons (IFN) and Tumor necrosis factor (TNF) signaling in endothelial cells after uptake^22^, whereas mitochondria phagocytosed by macrophages undergo fusion with the existing mitochondrial network and enhance respiration^23^. Morevoer, a recent study reported that adipocytes release damaged mitochondrial fragments into circulation in small EVs, which are then taken up by tissues including the heart and liver. A burst of ROS is generated in myocytes after uptake, which seems to precondition the heart against future stress^21^. Although some secreted EVs can function in cell-to-cell communication, our data suggest that secretion of EVs in response to lysosomal impairement functions primarily as an alternative quaity control mechanism to dispose of potentially harmful cargo. Ubiquitination is an important post-translational modification involved in targeting proteins and organelles for degradation^41^. In our study, both large and small EVs contained ubiquitinated cargo, and inhibiting EV release increased suceptibility to stress when internal degradation was compromised. Additionally, our observation that the phagocytosed EVs were delivered to lysosomes in macrophages is further evidence that this is an alternative disposal pathway.

In sum, our study delineates a novel process of mitochondrial quality control in cells when lysosomal-mediated degradation is compromised. The secretion of mitochondria in large EVs occurs via MVBs and involves inactivation of the small GTPase Rab7. As mitochondria are of bacterial origin, secretion of encapsulated mitochondria avoids activation of a potentially harmful inflammatory response. There is currently a strong interest in develping therapeutics that activates autophagy in the heart. However, increasing autophagosome formation to levels that exceeds the capacity of lysosomal degradation could lead to increased secretion of cargo by the cells. Although the impact of EV secretion on surrounding cells appears to be minimal, future studies need to elucidate whether excessive secretion affects the function of other cell types in the heart.

## Methods

### Animals

All animal experiments were performed following the Guidelines of National Institutes of Health on the Use of Laboratory Animals and approved by the Institutional Animal Care and Use Committee at the University of California, San Diego.

*Rab7^f/f^* mice^25^ were crossed with cardiac specific Myh6-MerCreMer (MCM) (Jackson Laboratory Stock #005650) to generate cardiomyocyte-specific *Rab7* knockout mice. Cre negative littermates were used as controls. To selectively delete *Rab7* in myocytes, male and female mice 8-10 weeks of age were injected (i.p.) with 40 mg/kg tamoxifen (Sigma-Aldrich, T5648) for five consecutive days. Control mice were also injected with tamoxifen. Cardiac-specific *Atg7* knockout mice^42^ were generated by crossing *Atg7^f/f^* mice with Myh6-Cre mice. Mice with cardiac specific expression of mito-Dendra2 were generated by breeding female PhAM-floxed^43^ (Jackson Laboratory, Stock #018385) and male Myh6-Cre or Myh6-MerCreMer mice. Lamp-2 knockout mice have previously been described^44^. Young (4-month-old) and Aged (24-month-old) male C57BL/6 mice were obtained from the National Institute of Health Institute of Aging colony (Charles River). For the in vivo chloroquine injection, saline or 80 mg/kg chloroquine (Sigma-Aldrich, C6628) was administered to mice by intraperitoneal injection, and the heart tissue were harvested 24 hours later analysis.

### Echocardiography

Transaortic echocardiography was performed using a Vevo 3100LT equipped with MX 400 transducer (30 MHz) for small animals as previously reported^45^. Mice were kept on a circulating water pad (37°C) and maintained under light anesthesia (0.5%–1% isoflurane, 98%–99.5% O2) during the procedure. M- and B-mode views were acquired, and cardiac function was analyzed using the VisualSonics software.

### Histological Analysis

To prepare paraffin sections, hearts were rinsed in PBS, fixed in 10% neutral buffered formalin for 48-72h and then transferred to 70% dehydrant alcohol for 48h. Hearts were embedded in paraffin, cut into 5-μm sections using a microtome (Leica Biosystems) and stained with hematoxylin and eosin (H&E) or Masson’s trichrome (Sigma-Aldrich). To prepare frozen sections, hearts were rinsed in PBS and then fixed in 10% neutral buffered formalin (Sigma) for 48-72h at 4 °C followed by incubation in 15% sucrose (6h) and 30% sucrose (12h) solutions. Hearts were frozen in Tissue-Tek^®^ O.C.T. Compound (SAKURA) for sectioning by a cryostat. For fluorescence staining, slides were incubated with primary antibody anti-CD68 (Invitrogen 14-0681-82) overnight. ImageJ was used to quantify fibrosis and CD68-positive cells.

### Mitochondrial respiration measurements

Mitochondria were isolated from *Rab7^f/f^* and *Rab7^f/f^ MCM* mouse hearts by investigators blinded to the genotype, and mitochondrial respiration was measured using the Seahorse XP Flux Analyzer (Seahorse Bioscience – Agilent Technologies) following a modified protocol described previously^46^. Data were analyzed using Wave for Desktop (Seahorse Bioscience – Agilent Technologies).

### Cell culture and Lentiviral transduction

WT and *Atg5*^-/-^ Mouse embryonic fibroblast (MEF) were obtained from Dr. Noboru Mizushima (The University of Tokyo, Japan). WT and *Rab7*^-/-^ MEFs were generously provided by Dr. Edinger (UC Irvine). Cells were cultured in Dulbecco’s Modified Eagle Medium (DMEM) containing 10% Fetal Bovine Serum (FBS), 1% Antibiotic-Antimycotic supplement at 37°C in 5% CO2. Raw 264.7 macrophages were cultured in RPIM 1640 containing 10% FBS, 1% Antibiotic-Antimycotic supplement at 37°C in 5% CO2. Cell lines were routinely tested for mycoplasma.

The Rab27 and Arl8b knockdown cell lines were established with the MISSION^®^ shRNA system. shRNA lentiviral transduction particles were purchased from Millipore-Sigma to deliver and stably express shRNAs in MEFs. MEFs were transduced with lentiviral particles for 48 h at 37 °C with polybrene (8 μg/ml) followed by puromycin (3μg/ml) selection for 48-72 h.

### Isolation of EVs

Large and small EVs were isolated using a modified differential centrifugation protocol as previously described^29^. Briefly, MEFs were seeded at density of 3 × 10^5^ cells/ml in a T182 flask and cultured in DMEM containing 10% EV-depleted FBS (Gibco) for 48h prior to collection of conditioned media. In some experiments, dimethyl sulfoxide (DMSO) or Bafilomycin A1 (5nM) were added to the cells after plating. The conditioned media was subjected to centrifugation at 300×g and 2,000xg (10 min each at 4°C) to remove unattached cells and large debris. The resulting supernatant was centrifuged at 10,000xg, 4°C for 50 min to pellet large EVs and the final supernatant was centrifuged in at 100,000xg for 70 min to pellet small EVs. Both large EV and small EV pellet were resuspended in Triton lysis buffer for Western Blot analysis or resuspended in PBS for nanoparticle tracking analysis (NTA) using NanoSight NS300.

To isolate EVs from mouse and human tissue, we utilized a previously published protocol^47^ with minor modifications. Briefly, heart tissues were cut into small pieces (2 × 2 × 2 mm) and then incubated for 45 min at 37°C either in Hanks’ Balanced Salt solution (HBSS) or plain DMEM media supplemented with Liberase HD (200 μg/ml, Roche) and DNase I (40 U/ml, Roche) with mild agitation (50rpm) to separate the EV from the cardiac tissue. After enzymatic digestion, dead cells and tissue debris were removed by filtration (70 μm filter) followed by successive centrifugation at 300 × g for 10 min and 2000 × g for 20 min. The resulting supernatant was filtered using 0.8μm filter unit (GE Healthcare) to eliminate apoptotic bodies and larger vesicles such as exophers^48^. The filtered supernatant was centrifuged at 16,500×g for 25 min and 118,000×g for 2.5 h to collect large and small vesicles, respectively. Large and small EV pellets were resuspended in Triton lysis buffer for Western blot analysis or resuspended in PBS for NTA using NanoSight NS300.

Alternatively, tissue and plasma EVs were isolated using an affinity purification method where the EVs are purified based on their surface markers (i.e. CD9, CD63, CD81) using the Miltenyi Biotec Pan Kit (human and mouse). Briefly, large EV- and small EV-enriched pellets obtained from differential centrifugation were resuspended in 500μl HBSS plus 50μL of Exosome Isolation MicroBeads (Miltenyi Biotec) recognizing cell surface markers CD9, CD63 and CD81, and incubated overnight at 4°C. Magnetic separation was performed and the EVs were eluted in Triton lysis buffer for WB analysis.

Mouse plasma samples were collected in K3 EDTA tube (Sarstedt Inc) and plasma were separated by centrifugation at 2000 × g for 15 min, followed by diluting with equal volume of PBS and centrifuged at 2000 × g for 30 min to remove cell debris and larger vesicles. The supernatant was incubated with MicroBeads (Miltenyi Biotec Pan Kit, mouse), then the magnetic separation and elution were performed on the next day.

Human heart biopsies were obtained at the University of California San Diego, Cardiology Department (Institutional Review Board approval #181206). Subjects gave their informed consent for use of their explanted cardiac tissues for research and that our study adheres to the principles of the declaration of Helsinki.

### EV uptake assay

Freshly isolated large EVs (10μg) from WT or *Rab7*^-/-^ MEF conditioned media were labeled with 5μM CFSE (Invitrogen) in PBS for 20 min at room temperature and kept protected from light, according to the manufacturer’s instructions. A control solution was prepared with CFSE in PBS in the absence of EVs. The staining reaction was terminated by adding 5ml cell culture medium plus 10% exosome depleted FBS. The CFSE-labeled large EVs were washed in PBS, centrifuged at 10,000xg for 50min and the resuspended in 20 μL of RPMI1640+10% exosome depleted FBS. The CFSE-labeled large EVS were added to RAW 264.7 macrophages (3× 10^4^ per 60mm MatTek dish) (MATTEK). After 20-24 h incubation at 37°C, LysoTracker Red (75nM) (Invitrogen) was added to the cells. After 0.5-1hr incubation at 37 °C, cells were fixed and imaged using Nikon Eclipse microscope. EV uptake rate was quantified as the number of CFSE-positive cells divided by total number of cells per image.

### Cell death assay

*Rab7*^-/-^ MEFs expressing control or Rab27 shRNA lentiviruses were treated with DMSO or 50 nM Bafilomycin A1(Millipore-sigma) or Oligomycin+ Antimycin A (10μM+10μM) (Sigma-Aldrich). To assess viability, cells were incubated with Yo-Pro1 (1:1,000; Life Technologies) and Hoechst 33342 (1:1000; Invitrogen) for 15 min at 37°C before imaging. Percent cell death was determined 24 h after treatments by dividing the number of Yo-Pro1-positive cells by total number of Hoechst 33342-positive cells as described^49^.

### Analysis of cardiac macrophages by flow cytometry

Adult mouse heart was excised, cannulated, and perfused with Liberase DH (26U/ml, Roche) in buffer containing 110 mM NaCl, 4.7 mM KCl, 0.6mM NaHPO4, 0.6 mM KH2PO4, 1.25 mM MgSO4, 10 mM KHCO3, 12 mM NaHCO3, 5.5 mM glucose, 30mM Taurine and 10 mM HEPES (pH 7.4). After digestion, ventricular tissue was gently teased into small pieces to dissociate loose cells. After about 30 min of sedimentation, the supernatant was centrifuged, and the cell pellet resuspended in staining buffer (BioLegend). Cells were seeded in 96-well plate, incubated with antibodies against CD45 (BioLegend, clone I3/2.3), CD11b (Tonbo Bioscience, clone M1/70) and F4/80 (Invitrogen, clone BM8), and then analyzed using a Guava benchtop mini-flow cytometer (EMD Millipore). Data were qualified using FlowJo software (Version 10).

### Real-time quantitative PCR

RNA was extracted from hearts using RNeasy Fibrous Tissue Mini Kit (Qiagen) and cDNA synthesized using the QuantiTect Reverse Transcription Kit (Qiagen). TaqMan primers *IL-6*, *Tnfα*, *Il-1b* were obtained from Thermo Fisher Scientific, while the TaqMan Universal Master Mix II was purchased from Applied Biosystems. The relative amounts of mRNA were normalized to *Rn18s* and the fold change of gene expression was calculated using the 2(-ΔΔCt) method. To measure mitochondrial DNA copy number in large EVs, genomic DNA was extracted using the GenElute Mammalian Genomic DNA Miniprep Kit (Sigma). *Rn18s* was used as a control for nuclear DNA content, and *cytochrome (Cyt) b* and *D-loop* were used for mtDNA quantitation as previously described^50^.

### Western blotting and antibodies

Proteins levels were analyzed by Western blotting followed our labs standard protocol as previously described^50^. Antibodies used in this study were: anti-Rab7 (9367S), anti-Alix (2171S), anti-CD81 (Mouse:10037S, human:56039S), anti-Tom20 (42406S), anti-Rab4 (2167S), anti-Rab5 (3547S), anti-Rab9 (5118S), anti-Rab11 (5589S), anti-LC3A/B (4108S), anti-Rab27 (69295S), anti-Arl8b (56085S) from Cell Signaling Technology; anti-SQSTM1/p62 (ab56416), anti-Tsg101 (ab83) from Abcam; anti-Tim23 (11123-1-AP) from Proteintech, anti-CD63 (PA5-92370) from Invitrogen; anti-Mitodendra2 (TA150090) from OriGene; anti-GAPDH (GTX627408) from GeneTex; and Anti-Ubiquitin (P4D1) (sc-8017) from Santa Cruz.

### Proteomics analysis

The proteomics analysis was done in collaboration with the Biomolecular/Proteomics Mass Spectrometry Facility at UCSD as described^51^. Briefly, proteins in isolated large and small EV fractions were separated by SDS–PAGE gel, subjected to in-gel tryptic digest and identified by ultra-high pressure liquid chromatography (UPLC) coupled with tandem mass spectroscopy (LC-MS/MS) using nano-spray ionization as described previously^50^. All plots were generated using the ggplot2 package in R. All Venn diagrams were generated using the VennDiagram package in R. Results were filtered by the number of unique peptide sequences detected. Proteins were used for analysis only if 2 or more unique peptides were detected. To account for low and missing values in the dataset, a background threshold value was calculated by taking the mean of all protein abundance values in the first quartile of the dataset, and this background value replaced missing values and all values lower than this threshold. Protein abundances were compared between large EV samples from WT and *Rab7*^-/-^ MEFs, and the −log10 adjusted p-values were plotted against the log2 fold changes. GO analysis was performed using the PANTHER statistical overrepresentation test (released 2022-02-02)^52^ on proteins significantly enriched in *Rab7*^-/-^ MEFs (t-test, p<0.05). The whole Mus musculus genome was used as background. Pathway overrepresentation was calculated using Fisher’s Exact test with a Benjamini-Hoechberg multiple testing correction for a false discovery rate of 5%. The top hits from GO biological process complete and cellular component complete were plotted according to −log10 false discovery rate. To compare our dataset to previously existing proteins found in different types of extracellular vesicles, data from Vesiclepedia (Version 4.1, 2018-08-15) and ExoCarta (2015-07-29) were downloaded and compared to the set of proteins common to all WT and *Rab7*^-/-^ MEFs. The DeepLoc2.0 web tool was used to predict subcellular localization of proteins in WT vs. *Rab7*^-/-^ MEFs.

### Immunofluorescence microscopy

MEFs were fixed with 4% PFA at 37°C for 15 mins and permeabilized with 0.2% Triton X-100 at 37°C for 30mins, followed by blocking in 5% normal goat serum and stained with primary antibodies (4 °C, overnight) as described previously^49^. To assess the colocalization between CD81 and mitochondria, WT MEFs were treated with Bafilomycin A1 (50nM) for 24h, and co-stained with anti-Cytochrome *c* (BD Biosciences, 556432) and anti-CD81 (Cell signaling, 10037S). To observe the co-localization between autophagosomes and EVs, both WT and *Rab7*^-/-^ MEFs were infected with an adenovirus expressing GFP-LC3 for 24 h prior to Bafilomycin A1 (50nM, 24 h) treatment. Cells were fixed and stained with anti-CD81(Cell signaling, 10037S). Fluorescence images were captured using a Nikon Eclipse microscope equipped with DS-Qi2 camera with 60X (oil immersion) objective. Colocalization was scored on a per cell basis and the software ImageJ-Fiji was used to calculate Pearson’s correlation coefficient (above the threshold).

### Transmission electron microscopy

Transmission electron microscopy was performed on heart sections from *Rab7^f/f^* and *Rab7^f/f^ MCM* mice as described previously^50^. At D28 post-tamoxifen administration, hearts were fixed in 2.5% glutaraldehyde in 0.1 M cacodylate buffer, and post-fixed in 1% osmium tetroxide. EV pellets were fixed with 2.5% glutaraldehyde, 4% paraformaldehyde in 0.1 M sodium cacodylate buffer pH 7.2 for 30 min at RT, rinsed in the same buffer 3x 10 min, followed by post-fixation in reduced 1% osmium tetroxide in buffer for 40 min, rinsed in the same buffer 3x 10 min, stained with 1% uranyl acetate for 40 min, rinsed with ultra-pure water for 3x 10 min, dehydrated in a graded acetone series until absolute (3x) for 10 min each step and finally embedded in Epon resin. Ultrathin sections of 50 nm were obtained with a diamond knife (Diatome) using a ultramicrotome UC7 (Leica), transferred to 300 mesh copper grids and stained with Uranyless (EMS) for 5 min. Sections were observed in a Philips CM100 transmission electron microscope (FEI) operated at 100 kV. For negative staining, isolated large and small EV pellets were resuspended in PBS and a 10 μL drop of each sample were absorbed on Formvar/Carbon coated grids. After 3 times of washing using water drop, the grids were stained 1 min with 2% Uranyl Acetate in water. Representative images were captured using a JEOL JEM-1400Plus transmission electron microscope (JEOL, Peabody, MA). Images were captured using a Gatan OneView digital camera (Gatan, Pleasanton, CA).

For the correlated light and electron microscopy (CLEM), cells were plated on gridded MatTek dishes (MatTek Corporation) and transfected with CD81-GFP (System Biosciences # CYTO124-PA-1) and mPlum-mito-3 (Addgene plasmid #55988) using Fugene6 (Promega Inc.) according to the manufacturer’s instructions. The images were captures as previously described^6^.

### Statistical analysis

All experiments were performed in the laboratory with a minimum of three independently biological replicates. Data are expressed as mean ± SEM. Statistical significance between groups were determined using one-way ANOVA or two-way ANOVA tests with Tukey’s post-hoc test or Student’s t-test.

## Acknowledgement

We would like to thank the University of California, San Diego - Cellular and Molecular Medicine Electron Microscopy Core (UCSD-CMM-EM Core, RRID:SCR_022039) for equipment access and technical assistance. The UCSD-CMM-EM Core is supported in part by the National Institutes of Health Award number S10OD023527. We also thank Dr. Siyi (May) Gu for her assistance with flow cytometry experiment and analysis.

Å.B. Gustafsson is supported by NIH grants R01HL155281 and R01HL157265. W. Liang is supported by the Tobacco Related Disease Research Program Postdoctoral Fellowship Award Grant T30FT0846. Illustrations were created with BioRender.com

## Contributions

ÅBG and WL designed the study, analyzed the experiments, and wrote the paper. WL performed and analyzed most of the experiments. RYD performed the proteomics analysis. BPW generated the Rab7 MCM mice and performed some of the initial characterization in cells and mice. LJL performed the mitochondrial respiration experiments. JMQ performed the CLEM experiment. EDA and JD provided the Lamp2 mice and human biopsies. DC and ADM assisted with the NTA experiments. RHN assisted with Western blot analyses and LC assisted with some of the mouse experiments. All authors reviewed the results and approved the manuscript.

## Conflicts

None

## Notes

### Competing Interest Statement

The authors have declared no competing interest.

