## Supplemental data for "The Small GTPase Rab7 Regulates Release of Mitochondria in Extracellular Vesicles in Response to Lysosomal Dysfunction"

### Supplementary Figures and Figure Legends

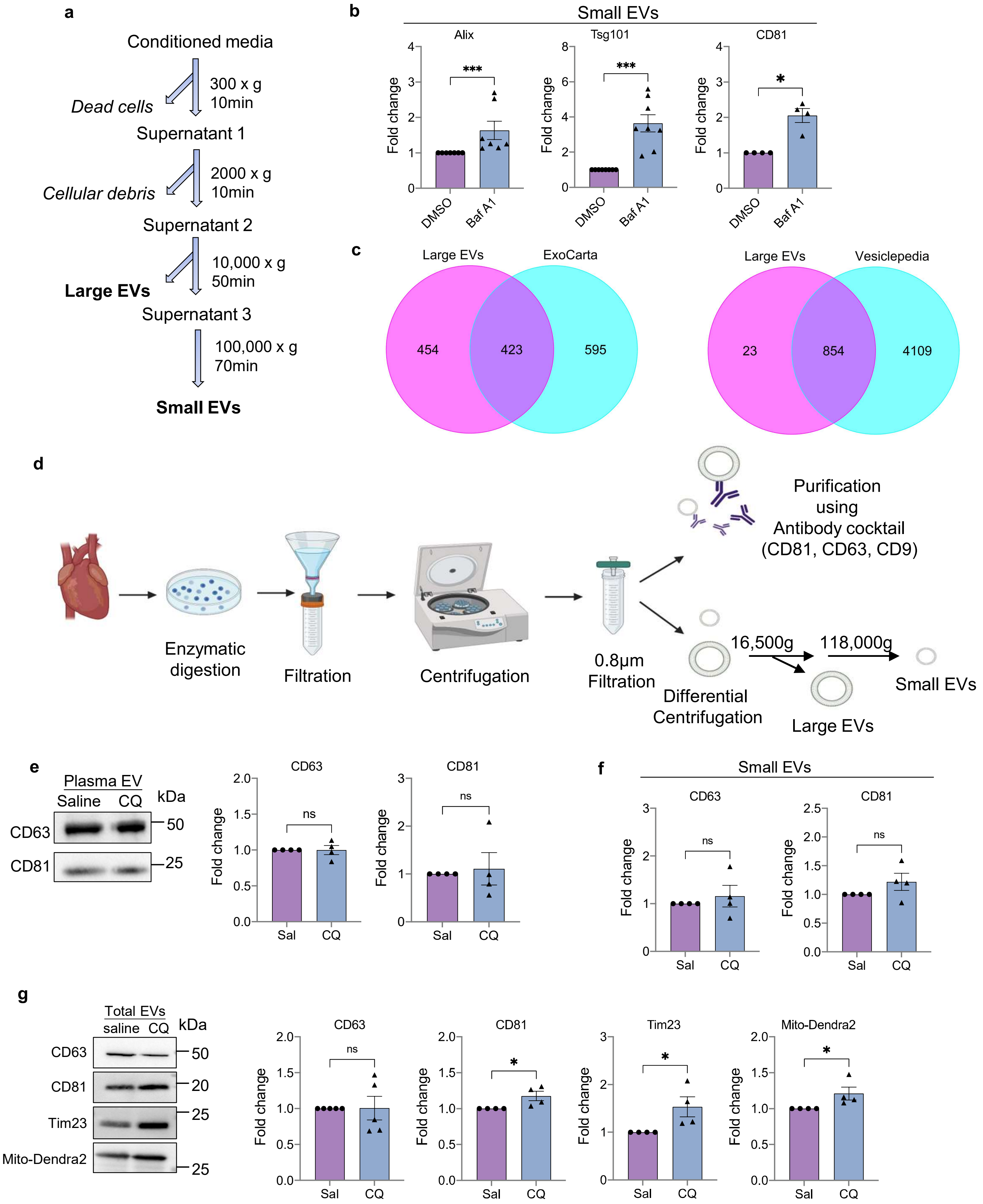

**Supplementary Figure 1. EV release in increased when lysosomal function is compromised. a.** Scheme of EV isolation from conditioned media using differential centrifugation. **b.** Quantification of proteins in small EV fraction from Fig. 1a (n=4-8). **c.** Venn diagrams showing the overlap of proteins identified in large EVs with those reported in Vesiclepedia and ExoCarta databases. **d.** Scheme of EV isolation from heart tissue. **e.** Western blot analysis and quantification of total EVs isolated from mouse plasma (n=4). **f.** Quantification of proteins in small EV fraction from Fig. 1d (n=4). **g.** Western blot analysis and quantification of CD63, CD81, Tim23, and Mito-Dendra2 protein levels in EV fractions isolated from heart tissue using immunoaffinity capture (n=4). Data are mean±SEM. *p<0.05, ***p<0.001, ns = not significant.

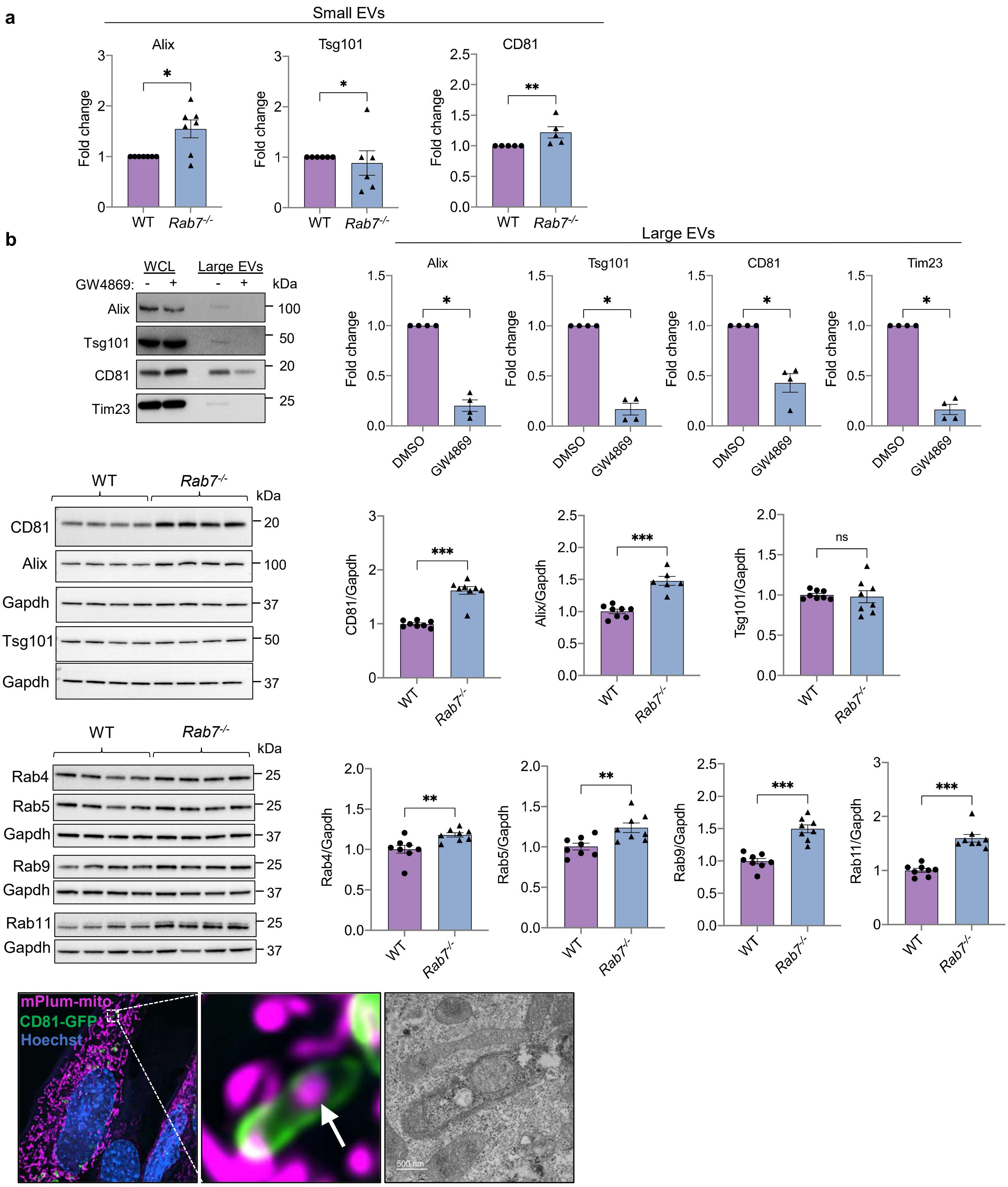
a
Small EVs
Alix
Tsg101
CD81
Fold change
WT
Rab7-/-
*
**
b
WCL
Large EVs
GW4869:
kDa
Alix
Tsg101
CD81
Tim23
100
50
20
25
Large EVs
DMSO
GW4869
c
WT
Rab7-/-
EV markers
CD81
Alix
Gapdh
Tsg101
37
CD81/Gapdh
Alix/Gapdh
Tsg101/Gapdh
***
ns
d
Endosomal proteins
Rab4
Rab5
Rab9
Rab11
Rab4/Gapdh
Rab5/Gapdh
Rab9/Gapdh
Rab11/Gapdh
e
mPlum-mito
CD81-GFP
Hoechst
500 nm

**Supplementary Figure 2. a.** Quantification of proteins in small EV fraction from Fig. 2a (n=5-7). **b**. Representative Western blot of proteins in whole cell lysates (WCL) and large EV fraction from *Rab7*$^{-/-}$ MEFs after treatment with GW4869 (2.5μM, 48hr). Quantification of proteins in the large EV fraction (n=4) **c**. Representative western blot and quantification of proteins in WT and *Rab7*$^{-/-}$ MEFs (n=6-8). **d**. Representative Western blot and quantification of proteins in WT and *Rab7*$^{-/-}$ MEFs (n=6-8). **e.** Correlative light and electron microscopy of *Rab7*$^{-/-}$ MEFs transfected with mPlum-mito3 and CD81-GFP. Fluorescent image and electron micrograph of a CD81-positive vesicle containing a mitochondrion are shown enlarged. Data are mean±SEM. *p<0.05, **p<0.01, ***p<0.001, ns = not significant.

**a**

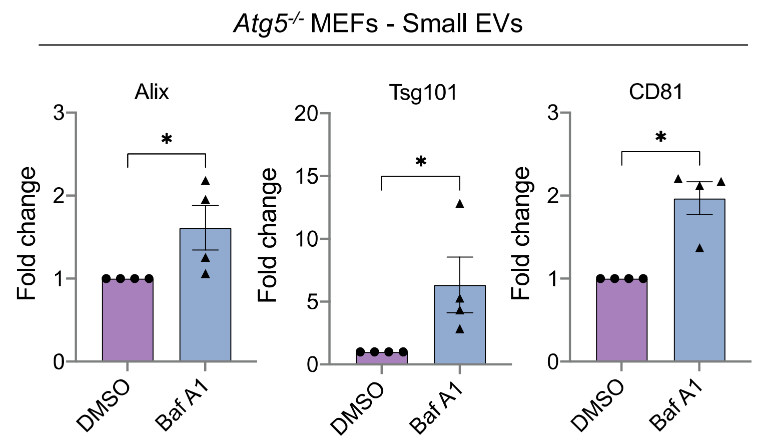


**b**

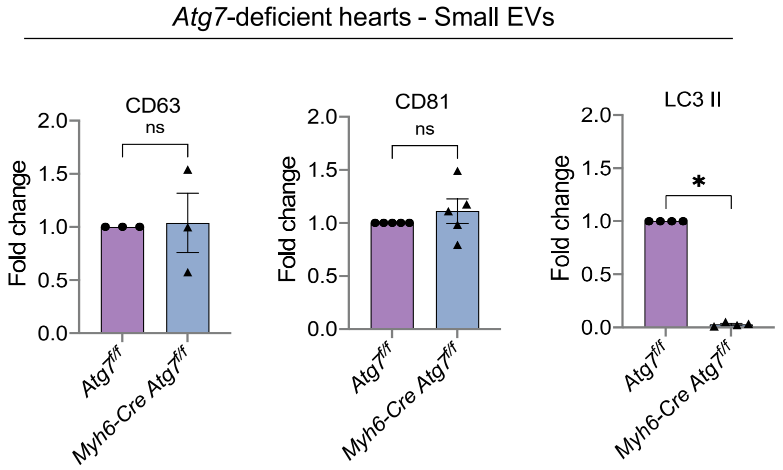


**Supplementary Figure 3. a.** Quantification of proteins in small EV fraction from Fig. 3a (n=4). **b.** Quantitation of proteins in small EV fraction from Fig. 3c (n=4-6) (n=4). Data are mean±SEM. *p<0.05, ns = not significant.

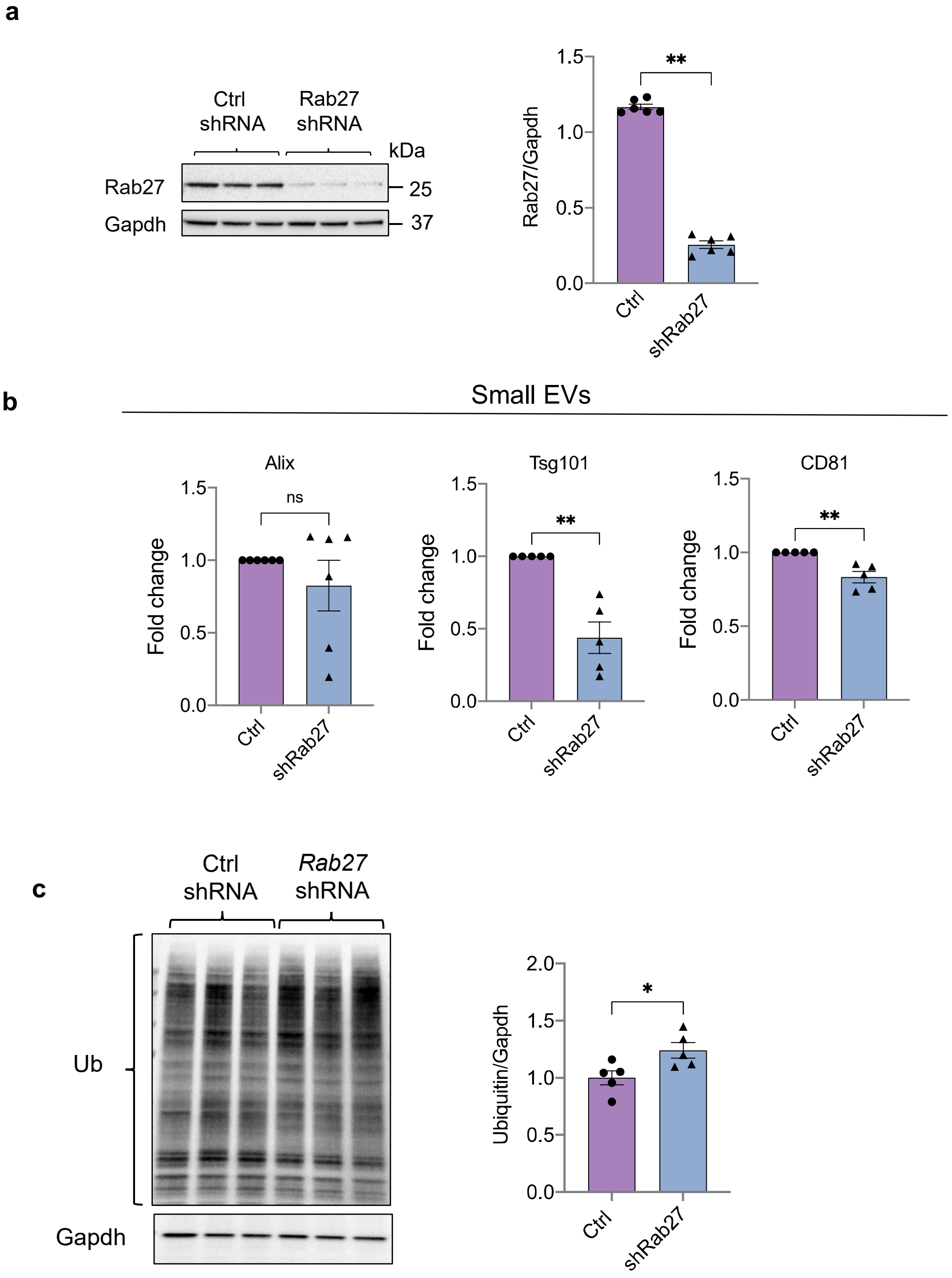


**Supplementary Figure 4. Effect of Rab27 knockdown in *Rab7*-deficient cells. a.** Confirmation of Rab27 knockdown in *Rab7*$^{-/-}$ MEFs. Representative western blot and quantification of Rab27 protein levels in WT and *Rab7*$^{-/-}$ MEFs after infection with control or Rab27a shRNA lentivirus (n=6). **b.** Quantification of proteins in small EV fractions from Fig. 4a (n=5-6). **c.** Western blot and quantification of ubiquitinated proteins in *Rab7*$^{-/-}$ MEFs after Rab27 knockdown (n=5-6). Data are mean±SEM. *p<0.05, **p<0.01, ns = not significant.

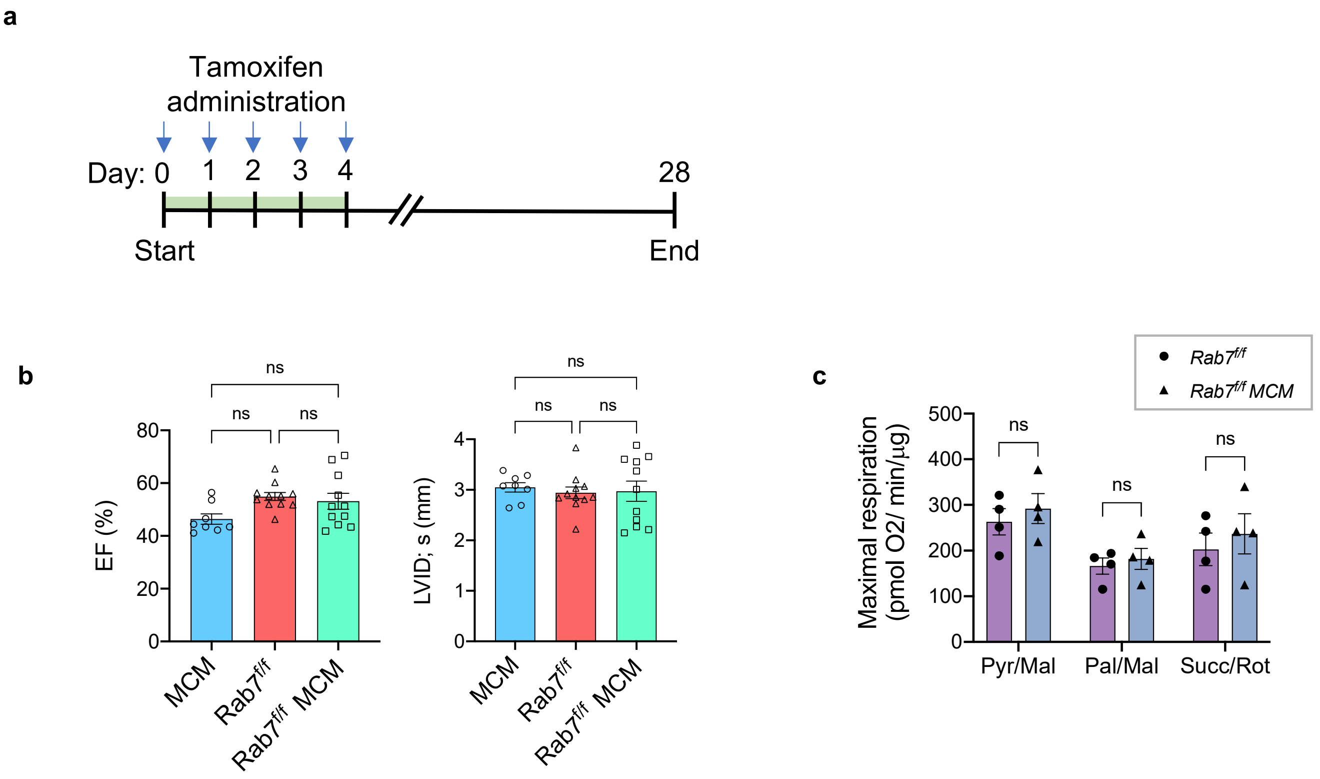


**Supplementary Figure 5. Characterization of *Rab7$^{f/f}$ MCM* mice. a.** Mice were injected with tamoxifen (40 mg/kg) for 5 days and all phenotyping conducted at 28 days post-tamoxifen treatment. **b.** Echocardiographic analysis of ventricular function and structure 28 days post-tamoxifen treatment. Percent (%) ejection fraction (EF) and left ventricular end-diastolic dimension in systole (LVED; s) . *MCM* (n=8), *Rab7$^{f/f}$* (n=11), *Rab7$^{f/f}$ MCM* (n=11). **c**. Assessment of mitochondrial respiration using isolated mitochondria from *Rab7$^{f/f}$ and Rab7$^{f/f}$ MCM* hearts show no differences in maximal respiration rates (FCCP uncoupled) with substrates for complex I (pyruvate/malate and palmitoyl carnitine/malate) or II (succinate/rotenone) (*n* = 4). Data are mean±SEM. ns = not significant.

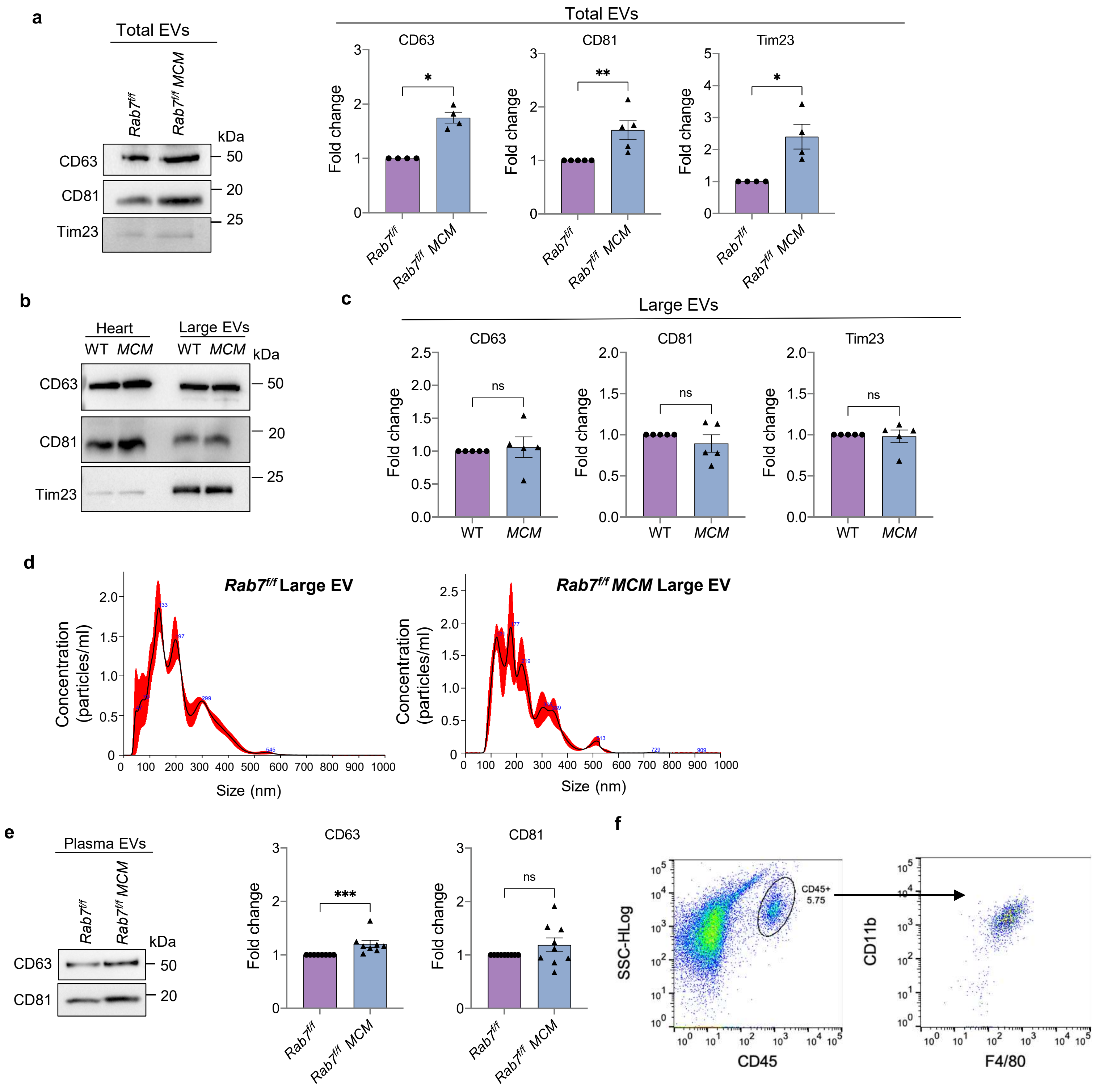


**Supplementary Figure 6. a.** Representative Western blot and quantification of proteins in large EV fractions isolated from hearts using immunoaffinity purification (n=8). **b.** Representative Western blot of protein levels in large EV fractions from WT and *MCM* heart tissue at D28 post-tamoxifen injection. **c.** Quantification of proteins in large EV fractions (n=5). **d.** Nano particle tracking analysis (NTA) to assess size distribution of large EVs isolated from *Rab7*$^{f/f}$ and *Rab7*$^{f/f}$ *MCM* heart tissue. **e.** Representative Western blot and quantification of proteins in total (large and small) EV fractions isolated from plasma using immunoaffinity purification (n=8). **f.** Gating strategy for the macrophage content analysis experiment shown in Fig. 6h. Data are mean±SEM. *p<0.05, **p<0.01, ***p<0.001, ns = not significant.

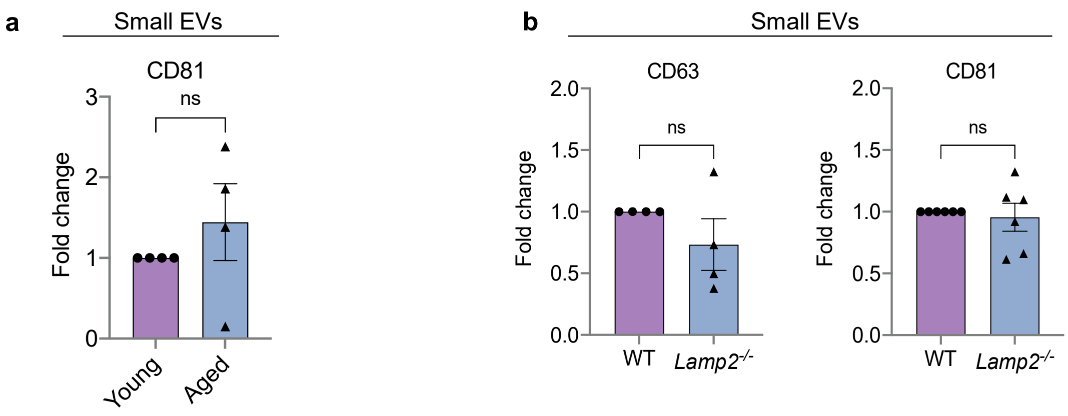


**Supplementary Figure 7. a.** Quantification of proteins in small EV fractions from Fig. 7a (n=4). **b**. Quantification of proteins in small EV fractions from Fig. 7e (n=4-6). Data are mean±SEM. ns=not significant.
